# Precision Functional Parcellation of the Human Cortex via Rest-Task fMRI Fusion

**DOI:** 10.64898/2026.06.11.731643

**Authors:** Da Zhi, Jingnan Du, Susan Whitfield-Gabrieli, Jörn Diedrichsen, Tian Ge

## Abstract

Individual-specific cortical parcellations enable the characterization of brain network organization that is often obscured by population-level atlases, with broad implications for both basic neuroscience and translational applications. However, existing methods rely primarily on resting-state fMRI and underutilize task-evoked data, which provide complementary information about functional specialization. This limitation partly reflects the challenge of integrating heterogeneous datasets that differ in task design, sample size, and cortical coverage. Here, we present mRBM-HBP, a scalable hierarchical Bayesian framework that incorporates a multinomial restricted Boltzmann machine to model spatial dependencies, enabling efficient and flexible integration of resting-state and task fMRI across diverse datasets and inference of both group-level and individual-level cortical parcellations. We show that mRBM-HBP achieves performance comparable to state-of-the-art resting-state-based parcellation methods while substantially reducing computational cost. By integrating large-scale task-fMRI datasets, we derive a task-based parcellation and demonstrate that resting-state and task conditions reveal largely consistent macroscopic networks, while task data provide state-specific refinements of functional boundaries. Moreover, a fused rest-task group-level atlas improves the accuracy, reliability, and individual specificity of inferred parcellations, particularly when individual-level data are limited. These results indicate that integrating resting-state and task fMRI enhances precision mapping of functional brain organization.

## Introduction

Over the past decade, advances in precision functional mapping (PFM) have substantially improved our understanding of inter-individual variability in brain organization by enabling reliable, individual-specific cortical parcellations (Gordon et al., 2017; Laumann et al., 2015). In contrast to traditional group-averaged approaches, PFM seeks to characterize each individual’s unique functional architecture, revealing network organization that is often obscured at the population level (Braga & Buckner, 2017; Gratton et al., 2018). This individualized perspective holds promise not only for basic neuroscience, but also for translational applications, including personalized diagnosis and prognosis, surgical planning, and identification of treatment targets in neurological and psychiatric disorders (Fox et al., 2012; Tie et al., 2014; Cole et al., 2020; Lynch et al., 2024; Sun et al., 2025).

Most existing individual-level parcellation methods rely on resting-state fMRI (Gordon et al., 2017; Kong et al., 2019, 2021), which captures intrinsic functional connectivity patterns of the brain (Biswal et al., 1995; Power et al., 2011; Yeo et al., 2011). In contrast, task-evoked fMRI provides complementary information by probing functional specialization through controlled engagement of specific cognitive processes (Duncan, 2010; Dosenbach et al., 2025). Emerging evidence suggests that functional organization shares core features across rest and task conditions but also exhibits task-dependent differences, reflecting both stable network architecture and context-dependent reconfiguration (Fair et al., 2007; Hasson et al., 2009; Cole et al., 2014; Krienen et al., 2014; Greene et al., 2020; Du et al., 2025; Zhi et al., 2025). Accordingly, applications involving the prediction of task-evoked responses or the delineation of functional boundaries may benefit from parcellations that incorporate task-relevant information (Glasser et al., 2016). However, task data have typically been used to validate whether parcel boundaries align with specific activation patterns, rather than being incorporated directly as inputs to parcellation models (Arslan et al., 2018).

Estimating parcellations under task conditions poses substantial methodological challenges. No single dataset currently provides both a sufficiently large sample size and a comprehensive set of tasks spanning all cognitive domains (Barch et al., 2013; Poldrack & Yarkoni, 2016; Eickhoff et al., 2018). Existing task-fMRI datasets are often limited in scope, focusing on specific domains or relatively small cohorts, which constrains their utility for learning generalizable functional organization (Eickhoff et al., 2018). Addressing this limitation requires methodological advances that can integrate information across heterogeneous datasets while accounting for differences in task design, sample characteristics, and data quality (Nettekoven et al., 2024; Zhi et al., 2025).

Here, we introduce a multinomial restricted Boltzmann machine (mRBM) to efficiently model spatial dependencies among brain locations and embed this architecture within a hierarchical Bayesian parcellation (HBP) framework (Zhi et al., 2025) for scalable inference in high-dimensional settings. The resulting mRBM-HBP model flexibly integrates resting-state and task fMRI data across multiple heterogeneous datasets to learn both group-level and individual-level parcellations. Using this model, we make two major contributions to precision functional mapping. First, we construct a task-based parcellation by integrating multiple task-fMRI datasets, showing that rest- and task-derived parcellations are highly similar while providing complementary information about functional organization. Second, to address a major bottleneck in PFM pipelines – the need for substantial amounts of resting-state data from each individual – we demonstrate that a fused rest-task group-level atlas improves the accuracy, reliability, and specificity of individual-level parcellations across independent datasets, particularly when individual-level data are limited. Together, these results establish task fMRI as a valuable complementary data source for advancing precision functional brain mapping.

## Results

### Overview of the mRBM-HBP model

Hierarchical Bayesian parcellation (HBP) is a Bayesian framework for the probabilistic integration of resting-state functional connectomes and task-evoked activation maps across diverse and heterogeneous datasets, enabling the estimation of both group-level and individual-level cortical parcellations. The framework comprises two key components (Figure 1a): (i) *Spatial arrangement model*, which defines a distribution over latent functional network labels at each cortical location and specifies how these labels are organized across the cortical surface. The group-level spatial arrangement model captures shared organizational structure across individuals and informs and regularizes individual-specific parcellations. (ii) *Dataset-specific emission models*, which define the likelihood of the observed data conditional on individual-level parcellations, thereby linking latent network assignments to modality- and dataset-specific functional measures. The parameters of both components are jointly estimated using an efficient Expectation-Maximization (EM) algorithm implemented through message passing and collaborative learning (Methods).

**Figure 1:**
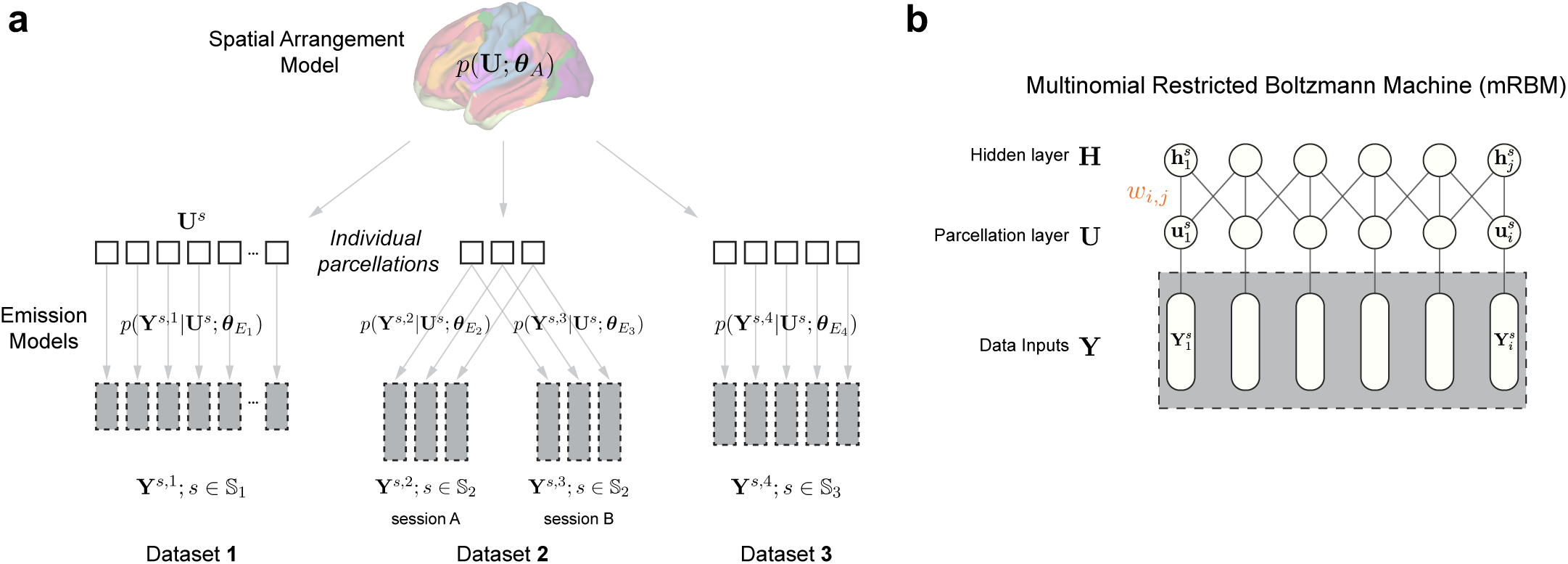
The mRBM-HBP model. **a**, Schematic of the Hierarchical Bayesian parcellation (HBP) framework with three training datasets. Observed data from individual *s* in dataset *n*, denoted **Y**^*s*,*n*^ for *s* ∈ 𝕊_*n*_, are represented by gray boxes, with box height indicating the amount of available data. Dataset 2 includes two sessions per individual. Hollow squares denote latent individual-level cortical parcellations, **U**^*s*^, which are informed and regularized by the group-level parcellation **U**. The group-level spatial arrangement model, *p*(**U**; *θ*_*A*_), defines a distribution over latent functional network labels at each cortical location, parameterized by *θ*_*A*_. Dataset-specific emission models, *p*(**Y**^*s*,*n*^|**U**^*s*^; *θ*_*E_n_*_), define the likelihood of the observed data conditional on individual-level parcellations, with parameters *θ*_*E_n_*_. **b,** Architecture of the multinomial Restricted Boltzmann Machine (mRBM) used as the spatial arrangement model. The mRBM comprises three layers: an input layer **Y**, a parcellation layer **U**, and a hidden layer **H**. For individual *s*, 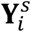 denotes the observed data (e.g., a resting-state functional connectivity profile or task-evoked activation profile) at cortical location *i*, while 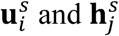 represent latent variables in the parcellation and hidden layers, respectively. The parcellation and hidden layers are interconnected by weights ω_*i,j*_, with no within-layer connections.

A fundamental challenge in neuroimaging data modeling is to capture spatial autocorrelation – the inherent correlation structure among nearby cortical vertices, which results in adjacent brain locations having a high probability of belonging to the same functional network. Existing Bayesian parcellation approaches use Markov random fields (MRFs) as spatial arrangement models, in which the probability of a latent network label at a given location depends on the labels of neighboring vertices, thereby encouraging spatial contiguity (Kong et al., 2019). However, MRF-based models are computationally expensive because posterior inference typically requires sequential, vertex-wise updates. To address this limitation, we developed a multinomial Restricted Boltzmann Machine (mRBM) as an alternative spatial arrangement model. In contrast to imposing explicit local dependencies among neighboring network labels, mRBM captures spatial correlations through a layer of hidden units that interact with the parcellation labels (Figure 1b). This bipartite architecture enables substantially more efficient model fitting in high-dimensional settings through parallel, layer-wise updates of network labels and connectivity weights (Methods).

During *model training*, mRBM-HBP leverages a corpus of datasets – comprising resting-state functional connectomes, task-evoked activation maps, or both, across multiple cohorts – to learn a group-level probabilistic parcellation by integrating all available information. Model hyperparameters, including the strength of the group-level prior and the degree of spatial smoothness, are tuned using an independent validation dataset. During *inference*, the learned group-level parcellation serves as a prior; given resting-state data from a new individual, mRBM-HBP infers an individual-specific probabilistic parcellation (Methods).

We performed extensive simulations to validate the mRBM-HBP framework using synthetic data generated from an MRF model with spatial dependencies among neighboring locations (Supplementary Methods; Supplementary Figure 1). These simulations demonstrated the benefit of incorporating a group-level prior for individual-level parcellation, relative to models trained solely on individual data, as well as the advantage of the mRBM model in capturing intrinsic spatial dependencies compared with approaches that do not explicitly model spatial smoothness (Supplementary Methods; Supplementary Figure 2).

### Benchmarking mRBM-HBP against resting-state parcellation methods in HCP

Most existing brain parcellation algorithms rely exclusively on resting-state fMRI. We therefore first evaluated the performance of mRBM-HBP when the group model was trained using resting-state data only. We benchmarked mRBM-HBP against MS-HBM (multi-session hierarchical Bayesian model), a widely used Bayesian cortical parcellation method that has demonstrated superior performance relative to alternative approaches (Kong et al., 2019). Analyses were conducted using data from the Human Connectome Project - Young Adult (HCP-YA) cohort. Each participant contributed approximately 60 minutes of resting-state fMRI data (4 runs × 15 mins), as well as task fMRI data spanning seven task domains, each acquired over two runs (Supplementary Tables 1-2). Group-level maps for both mRBM-HBP and MS-HBM were trained using all resting-state data from the same 40 individuals, with hyperparameters tuned in an independent set of 40 individuals. Each method generated a 17-network group-level parcellation (Methods).

Figure 2a-b show that the group-level parcellations generated by mRBM-HBP and MS-HBM were highly similar, with substantial Dice overlap across most network pairs (Figure 2c; Supplementary Table 3). Compared with the canonical Yeo-Krienen 17-network parcellation derived from a larger sample in the Harvard Brain Genomic Superstruct Project (Yeo et al., 2011), the overall network organization was also broadly consistent, with several noticeable differences (Supplementary Figure 3; Supplementary Table 4). For example, the mRBM-HBP HCP-YA parcellation separated the temporoparietal network into distinct language and auditory networks, introduced an additional visual network reflecting the visual hierarchy, and merged the two limbic networks into a single network (Supplementary Figure 3). Based on these differences, we updated the network labels to better reflect the organization observed in the HCP-YA dataset.

**Figure 2:**
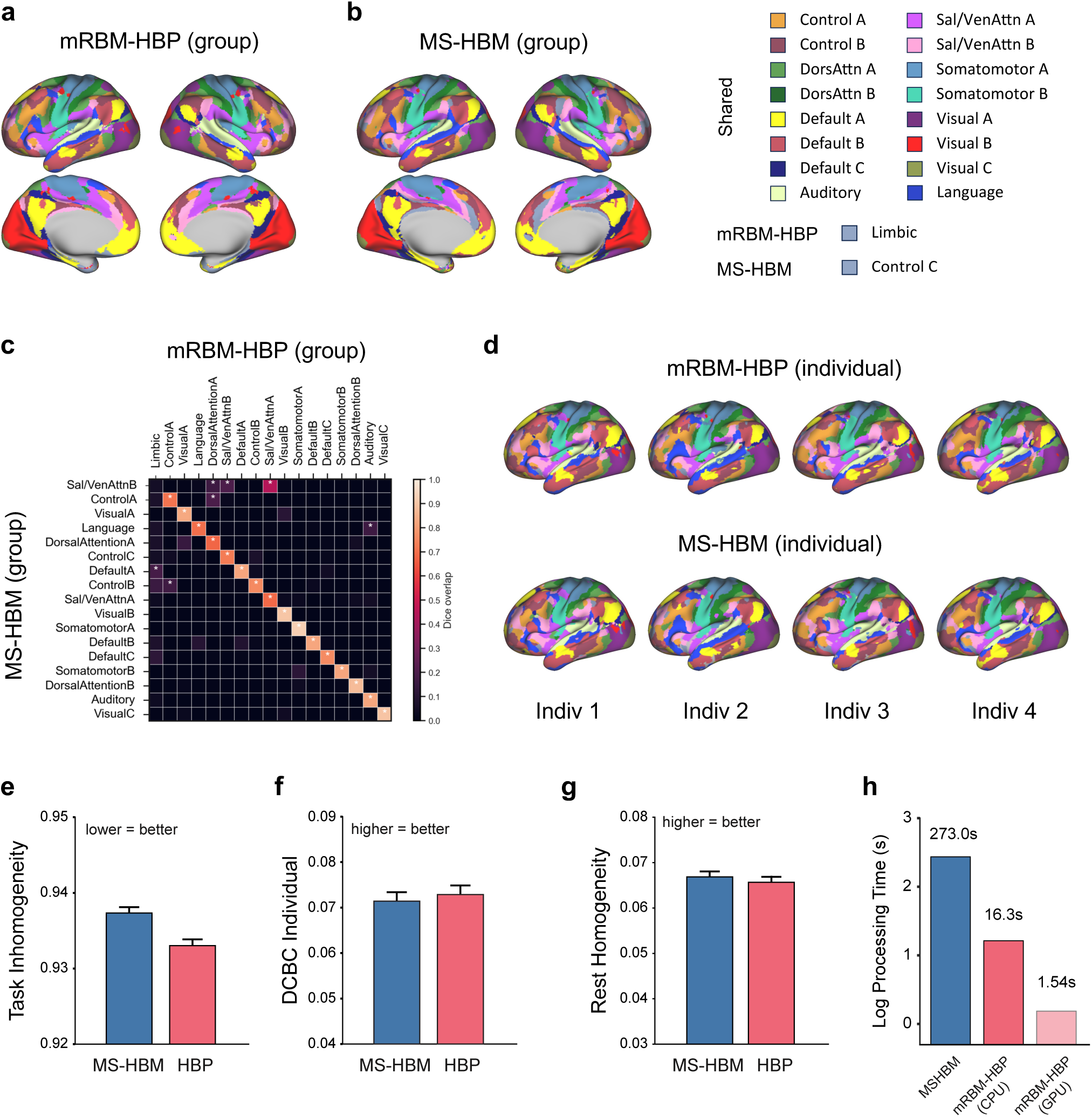
Benchmarking mRBM-HBP against MS-HBM using resting-state fMRI. **a-b**, 17-network group-level parcellations generated by mRBM-HBP and MS-HBM using resting-state fMRI data from 40 HCP-YA participants. Labels for four networks (Language, Auditory, Visual C, and Limbic) were updated from the canonical Yeo-Krienen 17-network parcellation to reflect the network organization observed in the HCP-YA dataset. **c,** Dice overlap between group-level parcellations generated by the two methods. **d,** Example individual-level parcellations from the HCP-YA test set generated by mRBM-HBP and MS-HBM using two resting-state runs per individual. **e-g,** Task inhomogeneity, DCBC (Distance-Controlled Boundary Coefficient), and resting-state homogeneity for individual-level parcellations generated by the two methods. Error bars represent the standard error of the mean. **h,** Average computational time required to generate parcellations for 10 individuals using MS-HBM, mRBM-HBP (CPU implementation), and mRBM-HBP (GPU acceleration).

Using these group-level parcellations, mRBM-HBP and MS-HBM were applied to generate individual-level parcellations for 200 randomly selected HCP-YA participants in the test set, independent of the training and validation sets, using two resting-state runs per individual. To evaluate performance under task conditions, we used 86 task-contrast maps spanning seven task domains from the same individuals (Supplementary Table 2) based on two complementary metrics: (i) task inhomogeneity, which quantifies the variance of task-evoked activation patterns across vertices within each network, averaged across networks and task conditions (Methods); and (ii) DCBC (Distance-Controlled Boundary Coefficient), which assesses boundary strength by comparing the functional similarity of vertex pairs within versus across network boundaries while controlling for spatial distance (Methods). To evaluate performance under resting-state conditions, we computed resting-state homogeneity, defined as the average correlation of fMRI time series among all pairs of vertices within each network, averaged across networks, using two held-out resting-state runs for each individual (Methods).

mRBM-HBP and MS-HBM produced visually similar individual-level parcellations (Figure 2d). However, when evaluated for their ability to predict functional boundaries under task conditions, mRBM-HBP achieved significantly lower task inhomogeneity (*p* = 1.56 × 10^−65^, paired *t*-test) and slightly higher DCBC than MS-HBM (Figure 2e-f; Supplementary Table 5). Under resting-state conditions, mRBM-HBP showed comparable, although slightly lower, resting-state homogeneity relative to MS-HBM (Figure 2g; Supplementary Table 6).

mRBM-HBP is also highly scalable and dramatically accelerates individual-level parcellation inference. Owing to the computational advantages of the mRBM architecture, which enables parallel updates of model parameters in contrast to the sequential MRF formulation used in MS-HBM, inference times were substantially reduced.

Specifically, MS-HBM required an average of 273.0 seconds to generate parcellations for 10 individuals, whereas mRBM-HBP required only 16.3 seconds using a standard CPU implementation and 1.54 seconds with GPU acceleration, representing approximately 16-fold and 180-fold speed-ups, respectively (Figure 2h).

Overall, these results demonstrate that, when trained and applied using resting-state fMRI, mRBM-HBP achieves performance broadly comparable to a state-of-the-art Bayesian cortical parcellation method, with improvements in task-based evaluation metrics, while substantially reducing computational cost.

### Task-based parcellation via integration of diverse task batteries across datasets

Having established the robustness of mRBM-HBP using resting-state fMRI, we next examined cortical network organization under task conditions. Task-based parcellation poses a greater methodological challenge because no single dataset provides a sufficiently comprehensive set of tasks spanning all functional domains at scale.

Effective task-based parcellation therefore requires integration of complementary task batteries across multiple datasets, a capability uniquely supported by the HBP framework (Zhi et al., 2025).

We trained a comprehensive task-based group atlas by leveraging three task-fMRI datasets that collectively span a broad range of task domains, including motor, visual, auditory, language, memory, executive function, emotion, social cognition, numerical processing, and higher-order cognition (Supplementary Table 1): (i) the Multi-Domain Task Battery (MDTB; number of participants, *N* = 24; number of unique task conditions, *N*_task_ = 47); (ii) the Nakai & Nishimoto dataset (*N* = 6; *N*_task_ = 103); and (iii) the Individual Brain Charting (IBC) dataset (*N* = 12; *N*_task_ = 208).

Figure 3a shows the task-based 17-network group-level cortical parcellation generated by mRBM-HBP. Despite being trained using different fMRI-derived features and a completely independent set of individuals, the resulting parcellation was remarkably similar to the group-level parcellation derived from HCP-YA resting-state fMRI (Figure 3b). Sixteen of the seventeen networks exhibited significant spatial overlap between the two parcellations (Figure 3c; Supplementary Table 7), with Dice coefficients ranging from 0.232 to 0.714. On average, unimodal networks showed greater overlap than heteromodal networks (Figure 3d). Even the network with the lowest Dice overlap, the Salience/Ventral Attention network (Dice = 0.232), exhibited highly similar cortical topology across the two parcellations (Figure 3e).

**Figure 3:**
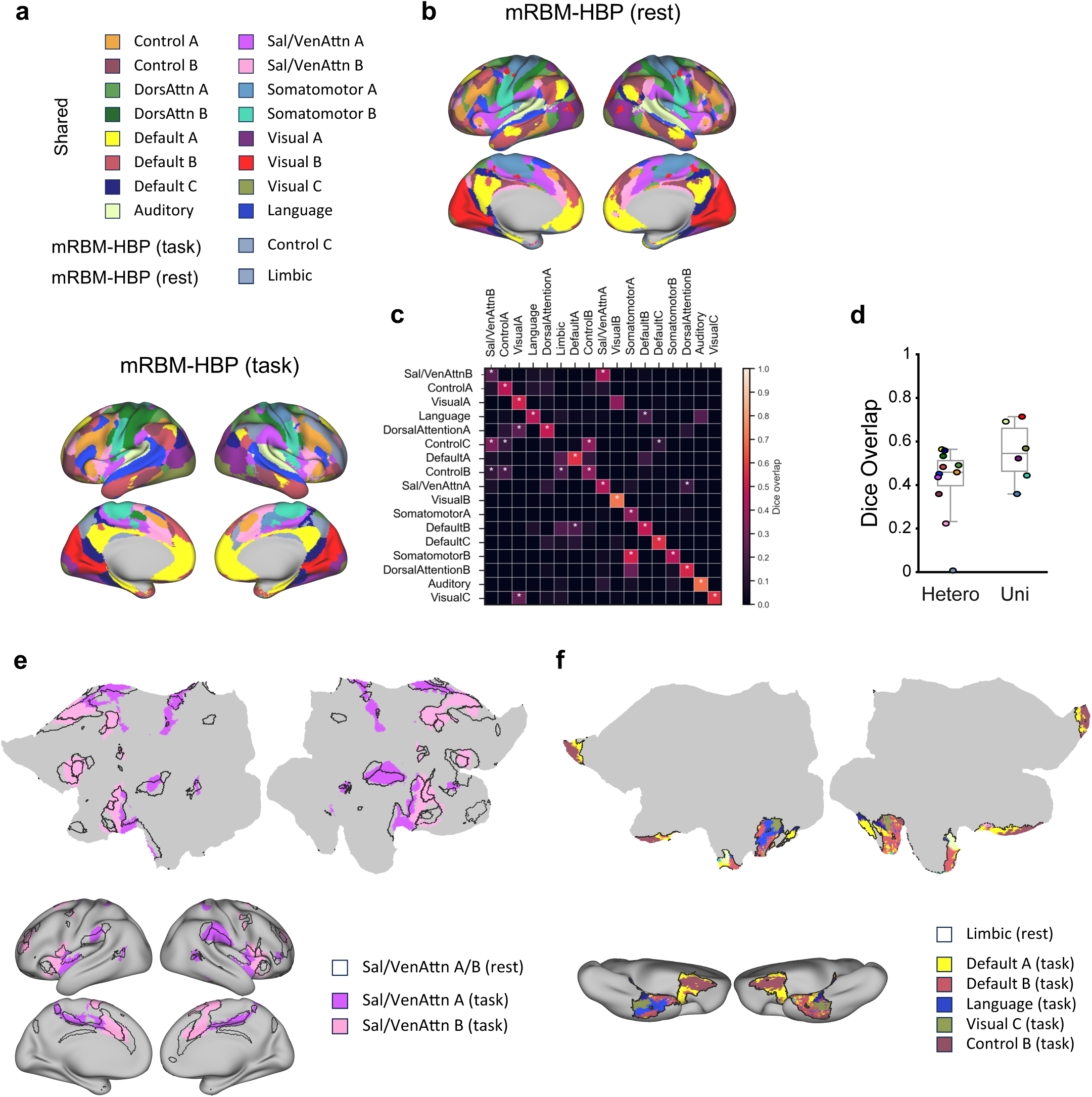
Comparison of resting-state and task-based 17-network group-level cortical parcellations. **a**, Task-based parcellation generated by mRBM-HBP by integrating three task-fMRI datasets spanning a broad range of functional domains. **b,** Resting-state parcellation generated by mRBM-HBP, reproduced from Figure 2a for comparison. **c,** Dice overlap between resting-state and task-based group-level parcellations. **d,** Distribution of Dice overlap for unimodal versus heteromodal networks. **e,** Comparison of the Salience/Ventral Attention networks between resting-state and task-based parcellations. **f,** Decomposition of the resting-state Limbic network into multiple networks under task conditions.

The only resting-state network lacking a clear task-based counterpart was the Limbic network, which decomposed into multiple networks, including default, language, control, and visual systems (Figure 3f). This pattern is expected: the Limbic network in resting-state parcellations occupies regions with low signal-to-noise ratio (Du et al., 2024), exhibits heterogeneous functional profiles, is prone to susceptibility artifacts, and is inconsistently reproduced across datasets and clustering methods. Regions traditionally assigned to the Limbic network at rest often participate in multiple distinct cognitive systems. In contrast, task activations provide stronger, more localized, and more functionally specific signals, thereby revealing finer-grained and domain-specific organization that is less apparent at rest. A detailed network-wise comparison between resting-state and task-based group-level parcellations is provided in Supplementary Figure 4.

Taken together, these results indicate that large-scale functional parcellations are largely consistent across resting-state and task conditions, while task fMRI provides complementary insights into the functional organization of the cortex.

### Rest-task fusion in a probabilistic group atlas improves individual-level parcellation in HCP-YA

Because resting-state and task-based parcellations capture both shared and complementary information, we hypothesized that integrating them into a fused rest-task group-level prior would improve individual-level parcellation. We therefore trained an mRBM-HBP model that jointly incorporated resting-state functional connectomes from the 40 HCP-YA training individuals and task activation maps from the MDTB, Nakai & Nishimoto, and IBC datasets, while balancing contributions from both modalities (Methods).

Figure 4a shows the fused 17-network group-level cortical parcellation, which integrated features from both resting-state and task-based parcellations. For example, the Limbic network observed in the resting-state parcellation was similarly decomposed into multiple networks, including Visual, Default, Language, and Control networks (Figure 3f). Conversely, the fused parcellation provided a clearer delineation of the Somatomotor A, comprising hand and upper-limb regions, and Somatomotor B, comprising face, tongue, and trunk regions, than the task-based parcellation (Figure 3a), a refinement primarily driven by information inherited from the resting-state parcellation (Figure 3b). Overall, the fused parcellation exhibited strong Dice overlap with both the resting-state parcellation (Figure 4b; Supplementary Table 8) and the task-based parcellation (Figure 4c; Supplementary Table 9). Network-wise probability maps of the fused group-level parcellation are shown in Supplementary Figure 5.

**Figure 4:**
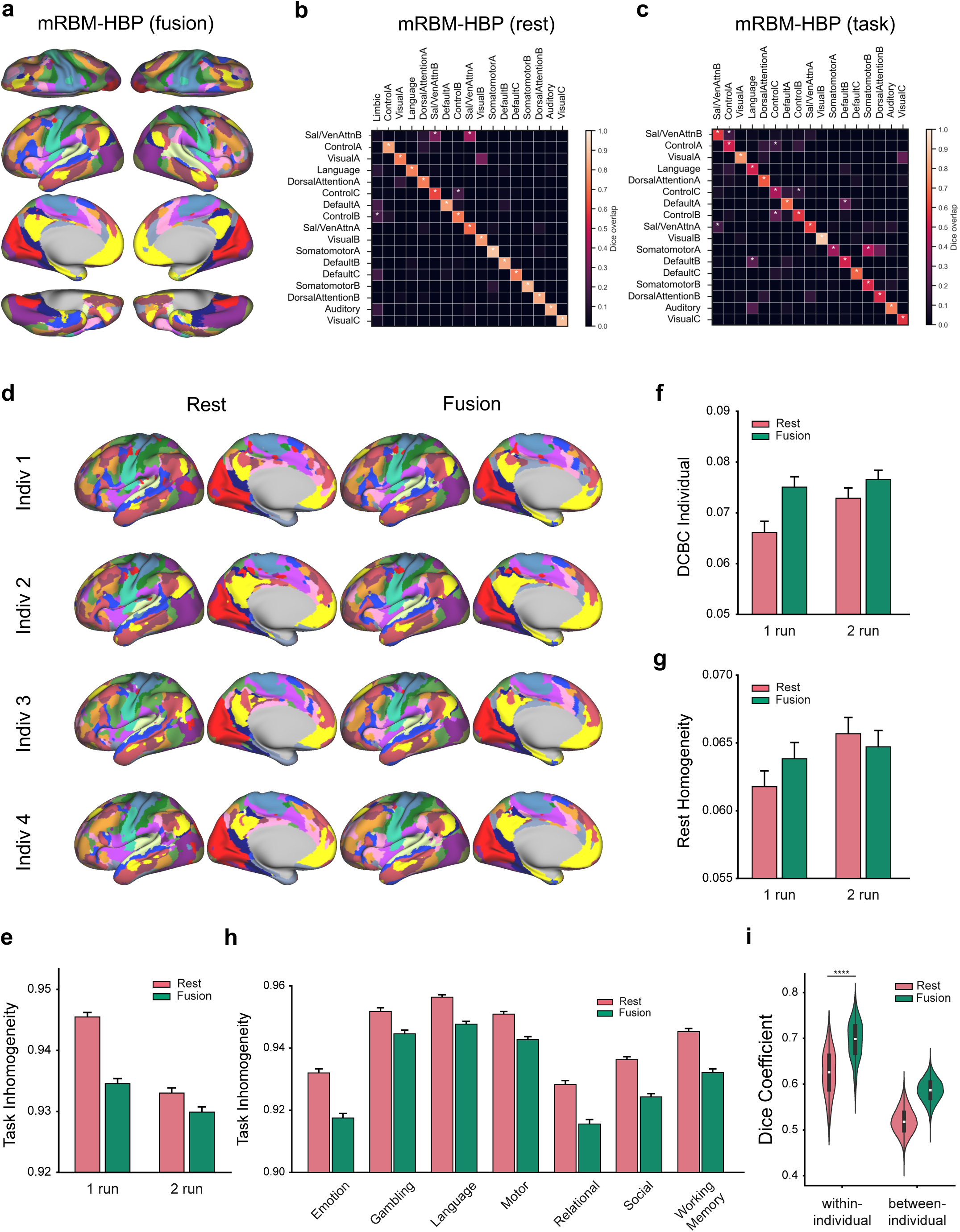
Fused rest-task group-level prior enhances individual-level parcellations relative to the resting-state prior. **a**, Fused rest-task 17-network group-level cortical parcellation generated by mRBM-HBP. **b,** Dice overlap between resting-state and fused group-level parcellations. **c,** Dice overlap between task-based and fused group-level parcellations. **d,** Example individual-level parcellations in the HCP-YA test set generated by mRBM-HBP using either the fused or resting-state group-level prior. **e-g,** Task inhomogeneity, DCBC (Distance-Controlled Boundary Coefficient), and resting-state homogeneity for individual-level parcellations inferred using either the fused or resting-state group-level prior, based on one or two resting-state runs per individual. Error bars represent the standard error of the mean. **h,** Task inhomogeneity across seven HCP-YA task domains for individual-level parcellations inferred using one resting-state run and either the fused or resting-state group-level prior. **i,** Within-individual reliability and between-individual similarity of cortical parcellations generated using the fused or resting-state group-level prior.

We then inferred individual-level parcellations for 200 participants in the HCP-YA test set using either the resting-state or fused group-level parcellation as the prior, based on one or two resting-state runs per individual. The resulting parcellations preserved group-level organization while capturing inter-individual variability (Figure 4d). For example, notable inter-individual differences were observed within default network regions in the medial precuneus and posterior cingulate cortex. We evaluated individual-level parcellations under both task and resting-state conditions, using 86 task-contrast maps spanning seven task domains and two held-out resting-state runs, respectively. As expected, longer resting-state scans (two runs) yielded more accurate parcellations regardless of the prior, as indicated by lower task inhomogeneity, higher DCBC, and higher resting-state homogeneity (Figure 4e-g; Supplementary Tables 5-6).

When comparing parcellations derived from the same amount of resting-state data, those informed by the fused prior consistently outperformed those based on the resting-state prior alone. Specifically, the fused prior yielded significantly lower task inhomogeneity (Figure 4e; 1 run: *p* = 1.12 × 10^−92^; 2 runs: *p* = 4.01 × 10^−90^; paired *t*-tests) and higher DCBC (Figure 4f; 1 run: *p* = 2.65 × 10^−23^; 2 runs: *p* = 8.20 × 10^−9^). Improvements were consistent across all task domains (Figure 4h; Supplementary Table 10). Notably, despite incorporating task information, the fused prior also improved resting-state homogeneity relative to the resting-state prior when using one resting-state run for inference (Figure 4g; *p* = 2.29 × 10^−51^), indicating enhanced parcellation quality across modalities. Across both task and resting-state evaluations, gains were more pronounced when only a single resting-state run was available (Figure 4e-g), suggesting that the fused prior provides the greatest benefit when individual-level data are limited.

These results demonstrate that integrating resting-state and task information into the group-level prior enhances the accuracy and robustness of individual-level parcellations in unseen individuals under both task and resting-state conditions.

### Fused rest-task prior improves within-individual reliability and individual identification

Lastly, we evaluated the within-individual reliability and between-individual variability of cortical parcellations generated by mRBM-HBP. For each individual in the HCP-YA test set, we inferred parcellations separately from the first and second resting-state runs (15 min each), using either the resting-state or fused group-level parcellation as the prior, yielding four parcellations per individual (Methods).

Visual inspection indicated that parcellations derived from independent runs of the same individual were highly consistent in the location, shape, and extent of functional networks, and were markedly more similar than parcellations from different individuals (Supplementary Figure 6). Quantitatively, for both priors, within-individual similarity, measured as the Dice coefficient between parcellations from the same individual, was substantially higher than between-individual similarity, defined as the Dice coefficient across all pairs of parcellations from different individuals (Figure 4i; resting-state: *p* = 1.23 × 10^−76^; fusion: *p* = 1.40 × 10^−90^, Welch’s *t*-test).

Comparing the two priors, parcellations informed by the fused prior exhibited significantly higher within-individual reliability than those generated using the resting-state prior (Figure 4i; average Dice coefficient: fused = 0.695 vs. resting-state = 0.624; *p* = 9.36 × 10^−128^, two-sample *t*-test; Supplementary Table 11). However, because between-individual similarity was also higher for the fused prior than for the resting-state prior (Figure 4i), higher test-retest reliability alone does not necessarily indicate improved individual specificity. We therefore performed an identification analysis in which, for each individual, the parcellation derived from run 2 was compared with all 200 parcellations derived from run 1, with identity assigned to the run-1 parcellation yielding the highest Dice similarity (Methods). We then repeated the analysis in the reverse direction. On average, the fused prior achieved higher identification accuracy than the resting-state prior across the 200 HCP-YA test individuals (95.5% vs. 90.7%; Supplementary Figure 6). Therefore, although the fused prior increased between-individual similarity (Figure 4i), it produced more robust and reliable within-individual parcellations while preserving individual-specific features, thereby enhancing their utility as functional “fingerprints” relative to parcellations derived from the resting-state prior alone.

### Comparison of resting-state and fused group-level priors in HPN

We replicated our analyses in the Harvard Precision Neuroimaging (HPN) dataset, which includes 15 right-handed adults aged 18-34 years (Du et al., 2024). Each participant was scanned across 8-11 sessions, with each session comprising multiple fMRI runs.

Across sessions, each participant contributed 17-24 resting-state runs (7 min 2 s per run) and an extensive set of task-based fMRI data spanning seven task domains: motor, visual, N-back, oddball, episodic projection, sentence processing, and theory-of-mind. The number of runs and scan durations varied across tasks (Supplementary Table 12). For the present study, we restricted analyses to 11 participants with complete task data (Methods).

A recent study in HPN demonstrated that a 15-network model is consistent with model-free seed-based analyses, exhibits high within-individual reliability, and captures key features of functional brain organization (Du et al., 2024). Motivated by this finding, we trained 15-network resting-state (Figure 5a) and fused group-level parcellations (Figure 5b; see Supplementary Figure 7 for network-wise probability maps) using the same 40 HCP-YA individuals and task activation maps from the MDTB, Nakai & Nishimoto, and IBC datasets.

**Figure 5:**
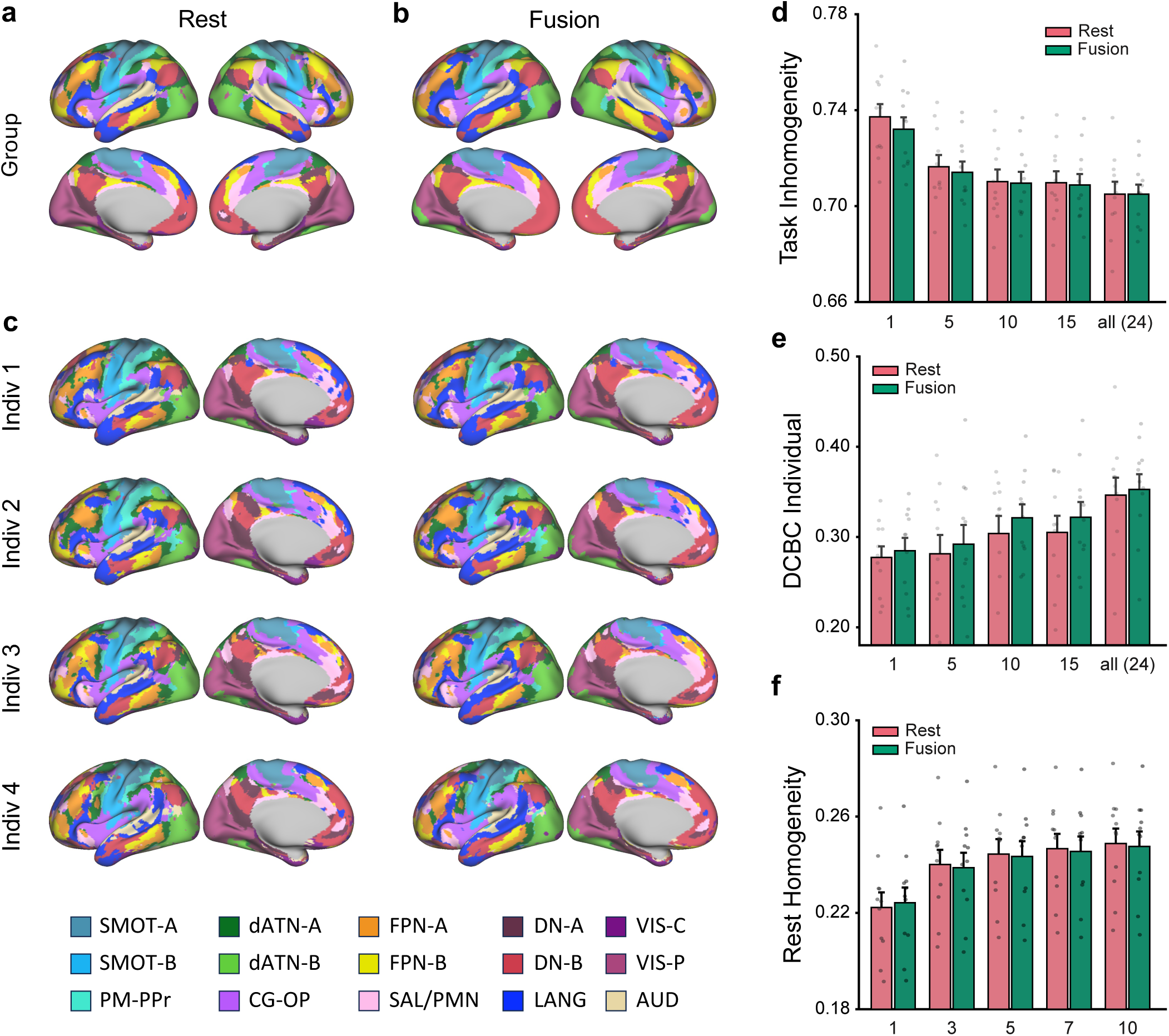
Comparison of resting-state and fused task-rest group-level priors in the Harvard Precision Neuroimaging (HPN) dataset. **a-b**, Resting-state and fused 15-network group-level cortical parcellations generated by mRBM-HBP. **c,** Example individual-level parcellations for HPN participants generated using all available resting-state runs. **d-f,** Task inhomogeneity, DCBC (Distance-Controlled Boundary Coefficient), and resting-state homogeneity for individual-level parcellations estimated using either the resting-state or fused group-level prior, based on varying numbers of resting-state runs. Error bars represent the standard error of the mean. SMOT-A, Somatomotor-A; SMOT-B, Somatomotor-B; PM-PPr, Premotor-Posterior Parietal Rostral; CG-OP, Cingulo-Opercular; SAL/PMN, Salience/Parietal Memory Network; dATN-A, Dorsal Attention-A; dATN-B, Dorsal Attention-B; FPN-A, Frontoparietal Network-A; FPN-B, Frontoparietal Network-B; DN-A, Default Network-A; DN-B, Default Network-B; LANG, Language; VIS-C, Visual Central; VIS-P, Visual Peripheral; AUD, Auditory.

Using these two group-level parcellations as priors, we then estimated individual-level parcellations for the 11 HPN participants (Figure 5c), varying the number of resting-state runs used for inference.

As expected, the accuracy of individual-level parcellations improved as more resting-state data were used, as reflected by lower task inhomogeneity and higher DCBC based on 17 task-contrast maps (Figure 5d-e; Supplementary Table 13), as well as higher resting-state homogeneity evaluated using five held-out runs (Figure 5f; Supplementary Table 14). Consistent with the HCP-YA test-set results, the fused prior consistently outperformed the resting-state prior in task-based evaluations, including task inhomogeneity and DCBC, when the same amount of resting-state data was used for inference (Figure 5d-e), while yielding comparable performance for resting-state homogeneity (Figure 5f). Performance gains tended to be more pronounced when individual-level data were limited (e.g., when only a single resting-state run was available), and diminished when all available resting-state runs were used for inference (Figure 5d-f), consistent with the group-level prior having less influence when abundant individual-level data are available.

### Fused rest-task prior improves parcellation accuracy under limited individual-level data in HPN

We applied mRBM-HBP independently to each resting-state run (7 min 2 s), generating multiple cortical parcellations per individual. Despite the limited data available in each run, network estimates were broadly consistent in their spatial organization across independent runs within the same individual (Figure 6a). For both the resting-state and fused priors, within-individual similarity was substantially higher than between-individual similarity (Figure 6b; Supplementary Table 15).

**Figure 6:**
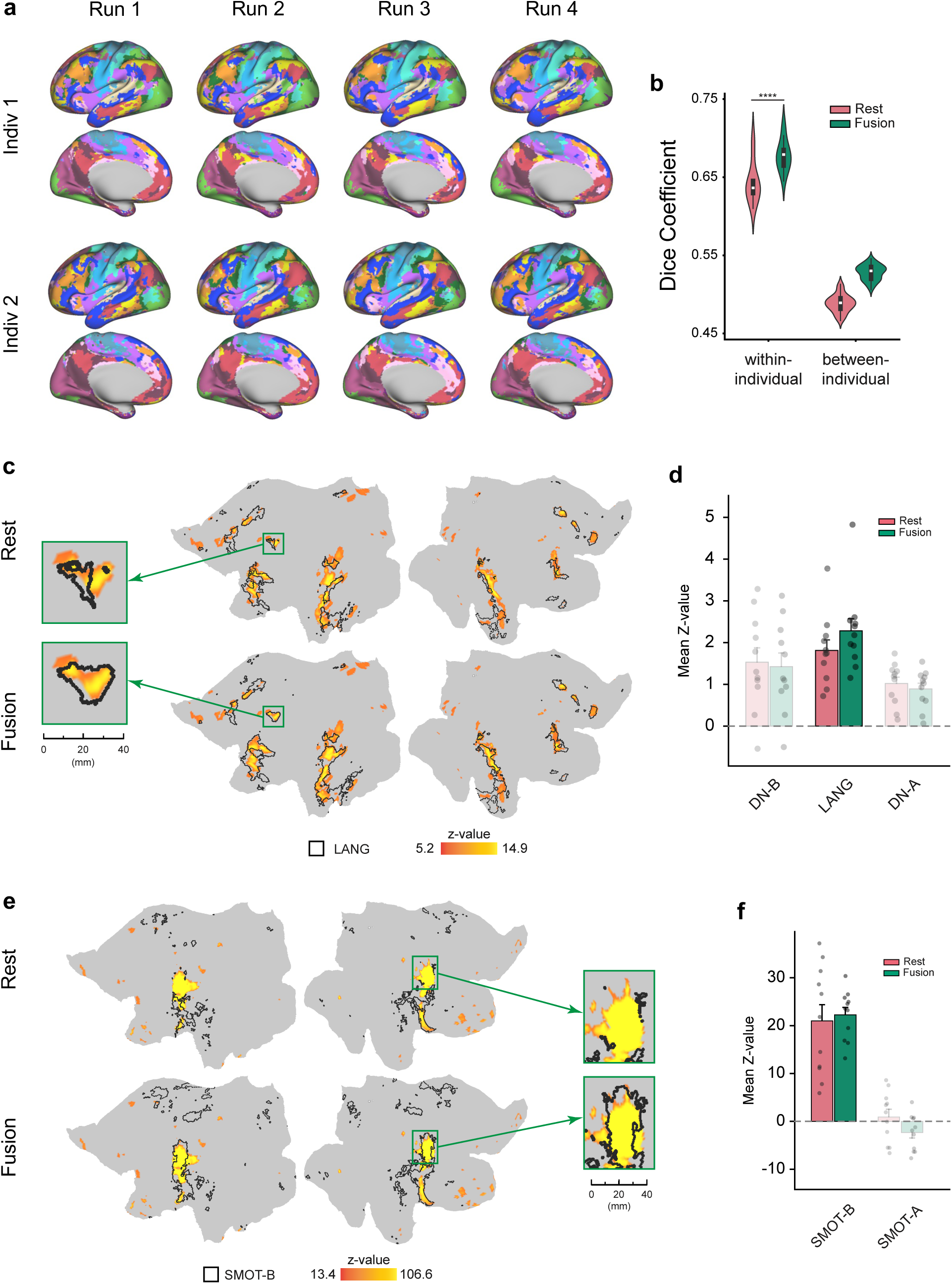
Reliability and precision of individual-level parcellations derived from mRBM-HBP in the Harvard Precision Neuroimaging (HPN) dataset. **a**, Example individual-level parcellations for two HPN participants generated using the fused prior from independent resting-state runs (7 min 2 s per run). **b,** Within-individual reliability and between-individual similarity of cortical parcellations generated using either the resting-state or fused group-level prior, based on a single resting-state run. **c,** Thresholded contrast map from the sentence processing task for an HPN participant, displayed on the flattened cortical surface and overlaid with Language network boundaries derived from the resting-state and fused priors using a single resting-state run. Zoom-in panels highlight activation in the left posterior middle frontal gyrus near area 55b. **d,** Average sentence processing contrast z-values within the Language network and adjacent default-mode subnetworks, DN-A and DN-B, defined using the resting-state and fused priors across HPN participants. **e,** Thresholded contrast map from the tongue movement task for an HPN participant, displayed on the flattened cortical surface and overlaid with Somatomotor-B network boundaries derived from the resting-state and fused priors using a single resting-state run. Zoom-in panels highlight activation in the right somatosensory cortex. **f,** Average tongue movement contrast z-values within the Somatomotor-A and Somatomotor-B networks defined using the resting-state and fused priors across HPN participants. In **d** and **f**, error bars represent the standard error of the mean.

Although improvements in task inhomogeneity and DCBC appeared modest in magnitude (Figure 5d-e), likely because these metrics summarize performance across tasks and networks, gains for specific tasks and their associated functional systems can be substantial. As an illustration, Figure 6c shows a thresholded contrast map from the sentence processing task for an HPN participant, displayed on the flattened cortical surface and overlaid with language network boundaries derived from the resting-state and fused priors using a single resting-state run. This task probes domain-specific processing of lexical and compositional semantics; contrasting sentences against nonword strings selectively activates the language network while differentiating it from adjacent systems, including auditory cortex and default-mode subnetworks (DN-A and DN-B).

Parcellations derived from the fused prior more precisely captured the spatial extent of language-related activation than those derived from the resting-state prior, particularly in the posterior middle frontal gyrus near area 55b in the left hemisphere – a region that is challenging to resolve at the individual level – as well as in the mid temporal cortex bilaterally. This improvement was further supported by higher average sentence processing contrast z-values within the Language network defined by the fused prior across HPN participants (Figure 6d; Supplementary Table 16).

Similar improvements were observed for the Somatomotor-B network, where parcellations derived from fused prior more accurately captured activation patterns associated with a tongue movement task than those derived from the resting-state prior (Figure 6e-f; Supplementary Table 17). Additional examples across other cognitive domains are provided (Supplementary Figure 8; Supplementary Tables 18-19). These results indicate that the fused prior provides more precise functional localization than the resting-state prior, particularly when individual-level resting-state data are limited.

In summary, the performance of mRBM-HBP and the fused rest-task group-level prior was robust across datasets, numbers of functional networks, task domains, and imaging acquisition protocols.

## Discussion

In this work, we developed mRBM-HBP, a hierarchical Bayesian parcellation (HBP) framework that incorporates a multinomial restricted Boltzmann machine (mRBM) to model the intrinsic spatial smoothness of neuroimaging data. Building on our prior work in cerebellar parcellation, this framework enables scalable and flexible integration of resting-state and task fMRI across diverse datasets and supports surface-based parcellation of the human neocortex at both group and individual levels. We show that mRBM-HBP achieves broadly comparable performance relative to existing methods under both resting-state and task conditions, while reducing computational cost by more than 100-fold with GPU acceleration. Leveraging this framework, we construct a task-based parcellation by integrating multiple task-fMRI datasets spanning a broad range of cognitive domains, demonstrating that rest- and task-derived parcellations are highly consistent yet provide complementary insights into functional organization. Furthermore, integrating resting-state and task data into a fused rest-task group-level atlas improves the accuracy, reliability, and specificity of individual-level parcellations, particularly when individual-level data are limited.

A key methodological advantage of this work is the explicit modeling of spatial dependencies among cortical vertices. Neuroimaging data exhibit intrinsic spatial smoothness, with neighboring cortical regions tending to share functional properties and engage in related cognitive processes (King et al., 2019; Kong et al., 2019; Zhi et al., 2022). Capturing this structure is essential for deriving stable and anatomically plausible parcellations, especially in the low-data regime typical of individual-level inference (Bijsterbosch et al., 2018; Amunts & Zilles, 2015; Schaefer et al., 2018). Although group-level parcellations naturally inherit smoothness through averaging across individuals, modeling spatial dependencies at the individual level remains challenging because high-resolution cortical mapping involves a massive number of vertices. Existing approaches either impose spatial priors within hierarchical Bayesian frameworks at substantial computational cost (Kong et al., 2019; Schaefer et al., 2018; Kong et al., 2021) or rely on assumptions of spatial independence (Zhi et al., 2025), limiting their ability to recover reliable, fine-grained functional topography.

We address this limitation by introducing mRBM as an alternative spatial arrangement model. The mRBM model captures spatial dependencies through a layer of hidden units that interact with parcellation labels via inter-layer connections across cortical locations. Our simulations demonstrated that mRBM-HBP reconstructs individual-level parcellations more accurately than spatially independent models, including those applied to smoothed data. In addition, in contrast to Markov random fields (MRFs), which typically require sequential, vertex-wise updates for posterior inference, the bipartite architecture of mRBM enables efficient parallel updates of network labels and model parameters, substantially reducing inference time. Unlike approaches that impose fixed or gradient-based spatial priors (Schaefer et al., 2018), mRBM does not assume a predefined form, scale, or structure of spatial dependencies, enabling data-driven learning of cortical organization. Together, this formulation promotes spatially coherent parcellations while avoiding overly restrictive priors that may bias network structure or obscure biologically meaningful variation.

A longstanding question in functional neuroimaging is whether brain organization observed during rest or task performance more accurately reflects underlying neurobiology (Smith et al., 2009; Cole et al., 2014; Krienen et al., 2014; Bijsterbosch et al., 2018). Over the past decade, resting-state fMRI has been the predominant approach for brain parcellation, owing to its ease of acquisition and the premise that spontaneous activity patterns capture intrinsic functional organization independent of specific task demands (Biswal et al., 1995).

Resting-state functional connectivity has been widely used to delineate stable large-scale networks and reproducible individual-level functional boundaries, forming the basis of many precision functional mapping efforts (Gordon et al., 2017; Kong et al., 2019; Schaefer et al., 2018; Yeo et al., 2011). In contrast, task-evoked fMRI characterizes brain organization under explicit functional demands, revealing networks engaged in specific cognitive, sensory, or motor processes (Dosenbach et al., 2025; Duncan, 2010; Fedorenko et al., 2010; Kanwisher et al., 1997; Saxe & Kanwisher, 2013). Although task data have historically been underutilized for parcellation due to variability across paradigms and limited data availability, they can reveal functional distinctions that may not be fully expressed at rest (Cole et al., 2014; Krienen et al., 2014; Eickhoff et al., 2018).

Emerging evidence suggests that functional boundaries estimated during task performance can systematically differ from those derived from resting-state data (Cole et al., 2014; King et al., 2019; Zhi et al., 2025), and may shift across cognitive states (Salehi et al., 2020). At the same time, major functional boundaries generalize across independent task contexts and are robustly detectable under both resting-state and task conditions (Smith et al., 2009; Cole et al., 2014; Tavor et al., 2016), indicating a shared underlying functional architecture. Recent work has suggested that individual-level parcellations inferred from dense, multi-domain task batteries may outperform those derived from resting-state data, and that functional boundaries become increasingly task-invariant as more task conditions are included in training (Nettekoven et al., 2026). However, acquiring hours of task-intensive fMRI is often infeasible in population cohorts or clinical settings. The present work provides a pragmatic alternative, demonstrating that integrating task data into a fused group-level prior significantly improves individual-level parcellations compared with a resting-state prior alone, while requiring only limited, accessible resting-state data from each individual.

Consistent with prior work, our results show that most task-derived networks exhibit one-to-one correspondence with resting-state networks, while specific networks show state-dependent differences, such as localized boundary shifts or topological reconfigurations, between task and rest. Together, these findings support a model in which core functional organization is largely preserved across brain states (Tavor et al., 2016; King et al., 2019), with context-dependent modulation in specific regions or systems. We posit that resting-state and task-based data emphasize different aspects of this shared architecture: resting-state data capture stable, intrinsic connectivity patterns, whereas task-evoked data provide targeted probes of functional specialization. This complementarity motivates methods that integrate both modalities to achieve more precise and functionally informative individual-level brain mapping (Cole et al., 2014; Tavor et al., 2016; Gratton et al., 2018).

Indeed, by integrating resting-state and task-evoked information into a fused group-level prior, we observed consistent improvements in individual-level parcellations across HCP-YA and HPN. Parcellations informed by the fused prior exhibited higher accuracy, greater within-individual reliability, and improved individual specificity compared with those based on a resting-state prior alone. Notably, these gains tended to be more pronounced when individual-level data were limited, a regime in which the group prior exerts greater influence. This setting is particularly relevant for real-world applications, including large-scale population studies, where extensive scanning is impractical, and clinical contexts, where prolonged imaging sessions are often infeasible (Kumar et al., 2024).

This work has several limitations and highlights directions for future research. First, constructing a task-based parcellation requires integrating task-fMRI datasets that collectively provide balanced coverage across diverse cognitive domains and sufficient activation coverage across the cortex. Over-representation of specific domains, cognitive systems, or cortical regions may bias the resulting parcellation. To mitigate this concern, we integrated three datasets – MDTB, Nakai & Nishimoto, and IBC – that together span a broad range of tasks without strong emphasis on any single domain. Nonetheless, residual biases or redundancy may remain, as multiple tasks can engage overlapping cognitive systems or cortical regions. We did not systematically assess how inclusion or exclusion of specific tasks or datasets affects parcellation quality. Future work should investigate optimal strategies for task selection and weighting, incorporate additional task-fMRI datasets that provide complementary information (Arafat et al., 2026), and determine the optimal balance between resting-state and task data when constructing fused parcellations.

Second, selecting the number of functional networks (*K*) remains a fundamental but unresolved challenge in brain parcellation and likely does not admit a single optimal solution (Eickhoff et al., 2018). Network-level parcellations typically use relatively small values of *K* (e.g., *K* < 50) to capture large-scale functional systems, whereas finer-grained parcellations may include hundreds of regions (Kong et al., 2021; Schaefer et al., 2018). The canonical 17-network model is widely used (Gordon et al., 2017; Yeo et al., 2011), although a recently proposed 15-network organization was shown to be highly reproducible at the individual level in densely sampled data and to align well with model-free seed-based estimates in HPN (Du et al., 2024). For task-based parcellations, the optimal *K* likely depends on the diversity and resolution of the task contrasts used for training. In this study, we constructed both 15- and 17-network parcellations to facilitate comparison with prior work and to enable rest-task fusion. The fact that mRBM-HBP and the fused rest-task prior yielded consistent advantages across both choices indicates that our conclusions are not contingent on a particular value of *K*. Nonetheless, developing principled, data-driven methods for selecting *K* and assessing the stability of network solutions remains an important direction for future research.

Third, we evaluated parcellation quality using complementary metrics under both task and resting-state conditions. However, each metric has inherent limitations, and no single measure provides a complete assessment of functional organization. Task inhomogeneity and resting-state homogeneity primarily assess within-network coherence and are relatively insensitive to boundary accuracy or between-network separation (Gordon et al., 2016). In contrast, DCBC evaluates local boundary sharpness but does not capture global homogeneity across the entire network and requires sufficiently rich task activation profiles for stable estimation. Metrics based on average activation (e.g., mean z-values) may favor parcellations that capture strongly activated regions while overlooking more spatially distributed but functionally relevant areas. In addition, aggregation performance across tasks and networks can obscure localized errors, region-specific misalignments, or domain-specific failures. We therefore interpret these metrics as capturing complementary aspects of parcellation quality. Future work should develop evaluation metrics that are more robust to preprocessing choices, spatial smoothness, and noise levels, and provide a more balanced assessment of the spatial localization, extent, and topology of functional patterns.

Lastly, although we integrated resting-state and task fMRI at the group level, individual-level inference in this study relied solely on resting-state data. The mRBM-HBP framework can be readily extended to incorporate individual-level task fMRI, which may further improve parcellation accuracy. Recent studies have also shown that functional networks estimated from task-residual time series closely resemble those derived from resting-state data (Cole et al., 2014; Du et al., 2025), suggesting the potential value of pooling resting-state and task-residual data (Du et al., 2025). Optimizing the design, acquisition, and integration of resting-state and task fMRI under realistic constraints, such as limited scan time or clinical feasibility, represents an important direction for advancing precision functional brain mapping.

## Methods

### Hierarchical Bayesian parcellation (HBP) framework for data fusion

HBP is a probabilistic framework for inferring both group-level and individual-level brain parcellations by integrating diverse and heterogeneous datasets. The framework consists of two key components (Figure 1a): (i) a *spatial arrangement model* and (ii) a set of *dataset-specific emission models*. For *P* cortical locations and a predefined number of networks *K*, the group-level spatial arrangement model, *p*(**U** ∈ ℝ^*K*×*P*^; *θ*_*A*_), defines a distribution over latent functional network labels, modeling the probability that each cortical location belongs to each functional network across individuals. This group-level model informs and regularizes individual-specific parcellations **U**^*s*^. Conditional on **U**^*s*^, the dataset-specific emission model, *p*(**Y**^*s*,*n*^|**U**^*s*^; *θ*_*E_n_*_), define the likelihood of the observed data **Y**^*s*,*n*^ for individual *s* in dataset *n*, thereby linking latent network assignments to modality- and dataset-specific functional measurements. Additional details of the model specification are provided in Supplementary Methods.

Model parameters for both components, {*θ*_*A*_, *θ*_*E_n_*_}, are jointly estimated using an efficient Expectation-Maximization (EM) algorithm implemented through a message-passing and collaborative-learning scheme (Supplementary Methods). The posterior expectations ⟨**U**⟩ and ⟨**U**^*s*^⟩ provide group-level and individual-level probabilistic parcellations, respectively, for the study sample 𝕊 = {𝕊_1_ ∪ 𝕊_2_ ∪ ⋯ ∪ 𝕊_*n*_}. Through this multi-level formulation, HBP enables flexible integration across datasets that differ in imaging modality, number of sessions, and signal-to-noise characteristics.

### The multinomial Restricted Boltzmann Machine (mRBM)

As a spatial arrangement model, mRBM – a multinomial extension of the original binary RBM (Hinton, 2002) – defines a distribution over latent functional network labels, 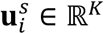, for each individual *s* and cortical location *i*. These labels represent the probability of assigning location *i* to each network, regularized by shared group-level parameters. The model comprises three layers (Figure 1b): (i) an input layer **Y**, containing observed imaging measurements across cortical locations, 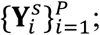 (ii) a parcellation layer **U**, containing latent network labels 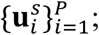 and (iii) a hidden layer **H** containing variables 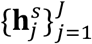. The input and parcellation layers are linked via a one-to-one mapping, whereas the parcellation and hidden layers are interconnected through connectivity weights ω_*i,j*_, with no within-layer connections.

Spatial dependencies across cortical locations are captured through interactions between the parcellation and hidden layers. Specifically, the connectivity weights ω_*i,j*_, modulated by a non-negative temperature parameter *θ*_w_, encode shared spatial structure across locations without requiring explicit pairwise potentials between neighboring vertices. In contrast to Markov random fields (MRFs), in which the probability of 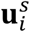 depends directly on neighboring labels 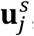, mRBM models spatial smoothness through its bipartite architecture. This design enables substantially more efficient inference in high-dimensional settings through parallel, layer-wise updates of network labels and connectivity weights (Supplementary Methods).

### Learning group- and individual-level parcellations using mRBM-HBP

The application of mRBM-HBP consists of two stages: (i) group-level model training and (ii) individual-level inference. During training, given a predefined number of functional networks, mRBM-HBP integrates information across individuals from multiple datasets and imaging modalities to learn a group-level probabilistic parcellation. Model hyperparameters, including the strength of the group-level prior and the degree of spatial smoothness, are tuned using an independent validation dataset. During inference, the learned group-level parcellation is used as a prior. Given resting-state data from new individuals, a dataset-specific emission model is fitted while keeping the parameters of the spatial arrangement model fixed. Individual-level parcellations are then obtained by computing the posterior expectations ⟨**U**^*s*^⟩, which combine the individual data likelihood with the group-level prior. Full details of model fitting and inference are provided in Supplementary Methods.

All model training and evaluation in this study were performed either on an NVIDIA RTX 6000 Ada Generation GPU with Python 3.10, CUDA 12.0, and PyTorch 2.5.1, or in a high-performance computing environment with GPU resources.

### Imaging datasets and preprocessing

We analyzed five datasets spanning resting-state and task-based fMRI across multiple functional domains (Supplementary Table 1). All datasets were either publicly available or provided by co-authors (Data Availability). All participants provided informed consent.

The *Human Connectome Project - Young Adult* (HCP-YA) dataset is a large-scale neuroimaging resource comprising high-quality structural, resting-state, and task fMRI data from healthy adults aged 22-35 years. Each participant contributed approximately 60 minutes of resting-state fMRI data (4 runs × 15 mins), preprocessed using the HCP minimal processing pipeline (Glasser et al., 2013). This yielded 1,200 time points per vertex in fsLR-32k surface space, with an initial spatial smoothing of 2 mm FWHM (full width at half maximum; Van Essen et al., 2012). We further applied a 4 mm FWHM Gaussian kernel, resulting in an effective smoothing of 6 mm FWHM. Participants also contributed task fMRI data spanning seven task domains, each acquired over two runs and processed using the HCP minimal processing pipeline (Glasser et al., 2013). For evaluation purposes, we used 86 task-contrast maps from the HCP S1200 release (Supplementary Table 2).

We analyzed three task-fMRI datasets that collectively span a broad range of functional domains: (i) the *Multi-Domain Task Battery* (MDTB; King et al., 2019); (ii) the *Nakai & Nishimoto* dataset (Nakai & Nishimoto, 2020); and (iii) a subset of the *Individual Brain Charting* dataset (IBC; Pinho et al., 2024). In the MDTB and Nakai & Nishimoto datasets, tasks were randomly intermixed within each imaging session, whereas in IBC, individual runs typically consisted of one or a small number of tasks within a specific cognitive domain. Task fMRI data from all three datasets were downloaded from open-access platforms (Data Availability) and preprocessed, as described in Zhi et al., 2025, using standard pipelines and resampled to fsLR-32k surface space. For each task condition, run-wise normalized task activation maps from first-level mass-univariate general linear model (GLM) analyses were aggregated across runs using the Functional Fusion toolbox (Code Availability). The resulting activation z-maps were further smoothed with a 6 mm FWHM Gaussian kernel using Connectome Workbench (Marcus et al., 2011), with appropriate handling of missing data, to improve the signal-to-noise ratio.

The *Harvard Precision Neuroimaging* (HPN) dataset is a densely sampled neuroimaging dataset in which 15 right-handed healthy adults aged 18-34 years underwent repeated MRI scanning, providing high-resolution resting-state and task fMRI for characterizing individual-specific brain organization (Du et al., 2024). Each participant was scanned across 8-11 sessions, with each session comprising multiple fMRI runs. Across sessions, participants contributed 17-24 resting-state runs (7 min 2 s per run), as well as an extensive set of task-based fMRI data spanning seven task domains, with varying numbers of runs per task (Supplementary Table 12). All fMRI data were acquired using a multi-EPI sequence and were preprocessed and projected to the fsaverage6 cortical surface with 2 mm FWHM smoothing using an in-house pipeline (Du et al., 2024). We then resampled the data to fsLR-32k space without additional smoothing. Task contrast maps for each task condition were generated following the procedures described in Du et al., 2024.

### Estimation of resting-state, task-based, and fused group-level atlases

The resting-state group-level parcellation was estimated using data from 40 HCP-YA participants matched to the training cohort in Kong et al., 2019. For each individual and each resting-state run, we computed Pearson correlations between the time series at each cortical vertex and the average time series within each of 642 regularly spaced hexagonal regions of interest (ROIs) per hemisphere. These ROIs were generated by projecting subdivided icosahedra from the sphere onto the fsLR-32k surface (Zhi et al., 2022). After excluding vertices in the medial wall and applying a hemispheric symmetry correction (Supplementary Methods), this procedure yielded a 59,518 × 1,210 functional connectome matrix. The matrix was then binarized by retaining the top 10% of correlations, following prior work (Kong et al., 2019; Yeo et al., 2011), and used as input features for the mRBM-HBP model.

The task-based group-level parcellation was estimated by integrating task activation maps across the MDTB (*N* = 24; *N*_task_ = 47), Nakai & Nishimoto (*N* = 6; *N*_task_ = 103), and IBC (*N* = 12; *N*_task_ = 208) datasets. For each individual within each dataset, each preprocessed activation map was thresholded to retain the original activation values for the top 10% most activated and bottom 10% most deactivated vertices. We verified that, across tasks, the selected vertices provided spatial coverage across the entire cortex (Supplementary Figure 9). We then stacked these task activation maps for each individual to form a vertices-by-tasks matrix, which served as input to the mRBM-HBP model.

The fused rest-task group-level parcellation was estimated by jointly integrating resting-state functional connectomes from the 40 HCP-YA training individuals and task activation maps from the MDTB, Nakai & Nishimoto, and IBC datasets using mRBM-HBP. During training, an internal weighting scheme was applied to balance contributions from resting-state and task data (Supplementary Methods). For all group-level analyses, we generated both 15-network and 17-network parcellations.

### Evaluation metrics for individual-level parcellations

We assessed the accuracy of individual-level parcellations under both task and resting-state conditions using complementary metrics.

*Task inhomogeneity* quantifies the variability of task-evoked activation within functional networks. Given a parcellation and a task-contrast map, we computed, for each network, the standard deviation of activation z-values across all vertices assigned to that network. These network-specific estimates were then averaged across networks, weighted by network size, to obtain an overall measure of task inhomogeneity (Gordon et al., 2017; Kong et al., 2019). Lower inhomogeneity values indicate more homogeneous task responses within networks, reflecting improved functional coherence and more accurate delineation of functional regions.

*Distance-Controlled Boundary Coefficient (DCBC)* quantifies the strength of functional boundaries defined by a parcellation by comparing the similarity of task-evoked activation profiles between vertex pairs within versus across network boundaries, while controlling for the intrinsic spatial smoothness of neuroimaging data (Zhi et al., 2022). Specifically, vertex pairs were grouped into bins according to their geodesic distance along the cortical surface. Within each distance bin, the similarity of activation z-value profiles across task contrasts was computed for within-network and between-network vertex pairs, and the difference between these similarities was evaluated. These differences were then averaged across distance bins, weighted by the reliability of estimates within each bin. Higher DCBC values indicate that the parcellation more accurately captures functional boundaries.

*Resting-state homogeneity* quantifies the functional coherence of a parcellation based on resting-state functional connectivity (Kong et al., 2019). For each network, homogeneity was computed as the average correlation of resting-state time series across all pairs of vertices within that network. These network-specific values were then averaged across networks, weighted by network size, to obtain an overall measure of resting-state homogeneity. Higher homogeneity values indicate greater similarity of intrinsic functional signals within networks, reflecting improved functional coherence and more accurate delineation of network boundaries.

*Mean z-values* were computed for run-averaged task contrasts within their corresponding target networks, as defined by each parcellation in the HPN dataset (Du et al., 2024). This measure quantifies the magnitude of task-evoked responses within the target network for each participant.

*Within-individual reliability* and *between-individual similarity* of cortical parcellations were quantified using the Dice coefficient (Dice, 1945). For each pair of parcellations, derived either from independent resting-state runs of the same individual or from different individuals, an iterative network-matching algorithm (Blumensath et al., 2013) was used to establish one-to-one correspondence between networks. The Dice coefficient was then computed for each matched network pair and averaged across networks to obtain an overall measure of similarity.

### Benchmarking mRBM-HBP against MS-HBM in HCP-YA

For mRBM-HBP, we used the 17-network group-level prior trained on 40 HCP-YA participants, each with four resting-state runs. Model hyperparameters, including the strength of the group-level prior and the degree of spatial smoothness, were tuned using an independent validation set of 40 HCP-YA participants. Individual-level parcellations were then inferred for 200 randomly selected HCP-YA participants, independent of the training and validation sets, using two resting-state runs per individual.

MS-HBM (multi-session hierarchical Bayesian model) was implemented following the instructions provided on the software Github page (Code Availability). We trained a 17-network group-level prior using MS-HBM on the same 40 HCP-YA participants, tuned the model on the same validation set, and inferred individual-level parcellations for the same 200 test participants using two resting-state runs per individual.

The resulting parcellations from both methods were evaluated under task conditions using 86 task-contrast maps spanning seven task domains (Supplementary Table 2), based on task inhomogeneity and DCBC. The released HCP-YA task contrast maps were used for evaluation without additional processing. Performance under resting-state conditions was assessed using resting-state homogeneity computed from two held-out runs of resting-state time series, with 2mm FWHM smoothing, for each individual.

### Comparing individual-level parcellations derived from resting-state and fused group-level priors in HCP

To compare the performance of resting-state and fused 17-network group-level parcellations for individual-level inference, both models were fine-tuned using an independent validation set of 40 HCP-YA participants. The models were then applied to infer individual-level parcellations for 200 test participants, using either one or two resting-state runs per individual. The resulting parcellations were evaluated under task conditions using the same 86 task-contrast maps spanning seven task domains (Supplementary Table 2), based on task inhomogeneity and DCBC. Performance under resting-state conditions was assessed using resting-state homogeneity computed from two held-out runs of resting-state time series, with 2mm FWHM smoothing, for each individual.

To assess reproducibility, for each group-level prior, two parcellations were inferred separately using independent resting-state runs from the same individual. Within-individual reliability was quantified as the Dice coefficient between parcellations derived from the same individual, whereas between-individual similarity was quantified as the Dice coefficient across all pairs of parcellations from different individuals.

To evaluate the extent to which parcellations capture individual-specific network organization, we performed an identification analysis. For each individual, the parcellation derived from run 2 was compared with all parcellations derived from run 1, and identity was assigned based on the highest Dice similarity. We repeated the analysis in the reverse direction, comparing run-1 parcellations against run-2 parcellations. Identification accuracy was computed by averaging results across both directions and all 200 HCP-YA test individuals.

### Replication in HPN

Analyses in the HPN dataset were conducted using a 15-network model and restricted to 11 participants with complete task data (Du et al. 2025). We first compared the performance of resting-state and fused group-level priors while varying the number of resting-state runs used for individual-level inference. The resulting parcellations were evaluated under task conditions using 17 task-contrast maps spanning seven cognitive domains (Supplementary Table 12), based on task inhomogeneity and DCBC. Performance under resting-state conditions was assessed using resting-state homogeneity computed from five held-out resting-state runs per individual.

To assess reproducibility under limited individual-level data, we independently inferred 15 parcellations per individual for each prior, each from a separate resting-state run. Within-individual reliability was quantified as the Dice coefficient across all pairs of parcellations from the same individual (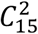^2^ = 105 pairs per individual), whereas between-individual similarity was quantified as the Dice coefficient across all pairs of parcellations from different individuals.

To evaluate functional specificity, thresholded contrast maps from selected tasks were visualized for representative HPN participants, with network boundaries derived from either the resting-state or fused group-level prior overlaid using matched resting-state runs. We then computed average task-contrast z-values within the corresponding target network across individuals.

### Data Availability

Raw fMRI data for the three task-based datasets are publicly available through OpenNeuro (https://openneuro.org/). The HCP S1200 release is publicly available through the Human Connectome Project Young Adult study website (https://www.humanconnectome.org/study/hcp-young-adult/document/1200-subjects-data-release). The HPN dataset is publicly available through the NIH Data Archive (https://nda.nih.gov/).

### Code Availability

The implementation of mRBM-HBP is publicly available at https://github.com/DiedrichsenLab/HierarchBayesParcel.

Code for preprocessing and integrating heterogeneous datasets is available at https://github.com/DiedrichsenLab/Functional_Fusion.

Code for the MS-HBM model and related data processing steps is available at https://github.com/ThomasYeoLab/CBIG/tree/master/stable_projects/brain_parcellation/Kong2019_MSHBM. The network correspondence toolbox (NCT) is available at https://github.com/rubykong/cbig_network_correspondence.

Connectome Workbench software (v1.5.0) is available at https://www.humanconnectome.org/software. Study-specific scripts, including those used to generate cortical functional probabilistic parcellations and conduct simulations, are available at https://github.com/dzhi1993/IndividualParcellation.

## Acknowledgements

The authors thank the investigators and participants of all neuroimaging datasets used in this study. Data were provided in part by the Human Connectome Project, WU-Minn Consortium (Principal Investigators: David Van Essen and Kamil Ugurbil; 1U54MH091657) funded by the 16 NIH Institutes and Centers that support the NIH Blueprint for Neuroscience Research; and by the McDonnell Center for Systems Neuroscience at Washington

University. The authors also thank Dr. Randy Buckner and colleagues for making the Harvard Precision Neuroimaging (HPN) dataset publicly available. T.G. is supported by National Institute of Mental Health (NIMH) R01MH130899.

