## Supplementary Information for "Precision Functional Parcellation of the Human Cortex via Rest-Task fMRI Fusion"

### Supplementary Methods

This supplementary material provides additional mathematical details of the mRBM-HBP model to support the Methods section in the main text.

For $P$ cortical locations and a predefined number of networks $K$, let $\mathbf{Y}$ denote the observed fMRI data (e.g., resting-state functional connectomes or task-evoked activation profiles), and let $\mathbf{U} \in \mathbb{R}^{K \times P}$ denote the matrix of network membership probabilities, where each column corresponds to a cortical location and each entry represents the probability of assignment to a given network. The mRBM-HBP model aims to infer the parcellation $\mathbf{U}$ by maximizing the joint probability:

$$p(\mathbf{Y}, \mathbf{U} | \boldsymbol{\theta}),$$

where $\boldsymbol{\theta}$ denotes the set of model parameters. Model learning proceeds by optimizing the expected log-likelihood $\mathcal{L} = \langle \log p(\mathbf{Y}, \mathbf{U}; \boldsymbol{\theta}) \rangle$ via variational inference (section S1). Under the HBP framework, $\mathcal{L}$ decomposes into two components: (1) the *spatial arrangement model* (section S2), which defines the expected arrangement log-likelihood $\mathcal{L}_A = \langle \log p(\mathbf{U}; \boldsymbol{\theta}_A) \rangle$, capturing the distribution of network assignment across cortical locations at the group level; and (2) the *dataset-specific emission models* (section S3), which define the data likelihood $\mathcal{L}_E = \langle \log p(\mathbf{Y} | \mathbf{U}; \boldsymbol{\theta}_E) \rangle$ for individuals from different datasets. The full parameter set $\boldsymbol{\theta}$ is thus partitioned as $\{\boldsymbol{\theta}_A, \boldsymbol{\theta}_E\}$ and estimated jointly using an EM-like algorithm described in section S4.

To model intrinsic spatial smoothness, we introduce a new spatial arrangement model based on a multinomial restricted Boltzmann machine (mRBM), which captures spatial dependencies among neighboring cortical locations on the surface. The (unnormalized) expected arrangement log-likelihood is given by:

$$\widetilde{\mathcal{L}}_A = \langle \boldsymbol{\eta} + \theta_w \cdot \mathbf{HW} \rangle,$$

where $\boldsymbol{\eta}$ is a matrix of log-probability parameters (see section S2), $\mathbf{H}$ denotes the hidden layer, $\mathbf{W}$ represents the connectivity weights between the hidden layer and parcellation layer $\mathbf{U}$, and $\theta_w$ is a temperature parameter controlling the overall connectivity strength. The mRBM model is trained using contrastive divergence under a stochastic maximum likelihood (SML) framework.

The mRBM-HBP model yields both group-level and individual-level parcellations. The group atlas is represented by network assignment probabilities at each cortical location $i$, defined as

$$p(\mathbf{u}_i) = \text{softmax}(\boldsymbol{\eta}_i),$$

which are assembled into the matrix $\mathbf{U}$. Individual-specific parcellations are obtained from the posterior distribution conditional on the data and model parameters:

$$p(\mathbf{U}^s | \mathbf{Y}^s; \boldsymbol{\theta}_A, \boldsymbol{\theta}_E) = \text{softmax}(\boldsymbol{\ell}^s + \boldsymbol{\eta} + \mathbf{H}^s \mathbf{W} \cdot \theta_w),$$

where $\boldsymbol{\ell}^s$ denotes the data likelihood of individual $s$ (see section S5). Detailed derivatives and implementation guidance are provided in sections S6-S8.

Section S9 describes the adjustment for left-right hemispheric symmetry based on the fs-LR32k surface mesh. Section S10 presents simulation studies of the mRBM-HBP model, including synthetic data generation (S10.1), the main results (S10.2), and two additional evaluation metrics for probabilistic parcellation: mean absolute error (S10.3) and mean adjusted expected cosine error (S10.4). We further show that the latter is mathematically equivalent to computing $1 - R^2$.

#### S1 Variational inference of the HBP framework

We use an EM-like algorithm to perform message passing and jointly learn the model parameters $\boldsymbol{\theta}$ within the HBP framework. We note that direct maximization of the marginal log-likelihood $\log p(\mathbf{Y}^s; \boldsymbol{\theta})$ is intractable, as it requires summing over all possible configurations of the latent variables (here, the individual-specific parcellations $\mathbf{U}^s$). To address this, we introduce a variational distribution $q(\mathbf{U})$ over the latent variables and optimize the evidence lower bound (ELBO) (Wainwright and Jordan, 2008; Blei et al., 2017). Let $\mathbf{Y}$ denote the collection of all datasets. For notational simplicity, consider a single individual $s$. The ELBO is derived as:

$$\begin{aligned}
\log p(\mathbf{Y}|\theta) &= \log \sum_{\mathbf{U}} p(\mathbf{Y}, \mathbf{U}|\theta) \\
&= \log \sum_{\mathbf{U}} q(\mathbf{U}) \frac{p(\mathbf{Y}, \mathbf{U}|\theta)}{q(\mathbf{U})} \\
&\geqslant \sum_{\mathbf{U}} q(\mathbf{U}) \log \frac{p(\mathbf{Y}, \mathbf{U}|\theta)}{q(\mathbf{U})} && \text{(Jensen's inequality)} \\
&= \langle \log p(\mathbf{Y}, \mathbf{U}|\theta) - \log q(\mathbf{U}) \rangle_q \triangleq \mathcal{L}(q, \theta) - \log \langle q(\mathbf{U}) \rangle_q && (1)
\end{aligned}$$

This provides a lower bound on the marginal log-likelihood.

Summing over all individuals $s$ and datasets $n$, we obtain:

$$\sum_{s,n} \log p(\mathbf{Y}^{s,n}; \boldsymbol{\theta}) \geqslant \sum_{s,n} \langle \log p(\mathbf{Y}^{s,n}, \mathbf{U}^s; \boldsymbol{\theta}) \rangle_q - \langle \log q(\mathbf{U}^s) \rangle_q. \tag{2}$$

The first term corresponds to the expected joint log-likelihood $\mathcal{L}$, which decomposes into emission and arrangement components:

$$\begin{aligned}
\mathcal{L} = \sum_{s,n} \langle \log p(\mathbf{Y}^{s,n}, \mathbf{U}^s; \boldsymbol{\theta}) \rangle_q &= \sum_{s \in \mathcal{S}_1} \langle \log p(\mathbf{Y}^{s,1} | \mathbf{U}^s; \boldsymbol{\theta}_{E1}) \rangle_q + \sum_{s \in \mathcal{S}_2} \langle \log p(\mathbf{Y}^{s,2} | \mathbf{U}^s; \boldsymbol{\theta}_{E2}) \rangle_q \\
&+ \ldots + \sum_{s} \langle \log p(\mathbf{U}^s; \boldsymbol{\theta}_A) \rangle_q \triangleq \mathcal{L}_{E1} + \mathcal{L}_{E2} + \ldots + \mathcal{L}_A, && (3)
\end{aligned}$$

where $\boldsymbol{\theta}_A$ denotes parameters of the spatial arrangement model, and $\{\boldsymbol{\theta}_{E1}, \boldsymbol{\theta}_{E2}, \ldots\}$ denote parameters of dataset-specific emission models. This partition enables separate updates of the arrangement and emission models.

In the E-step, the variational distribution is updated to approximate the posterior given the current estimates of model parameters:

$$\begin{aligned}
q(\mathbf{U}^s) &= p(\mathbf{U}^s | \mathbf{Y}^{s,1}, \mathbf{Y}^{s,2}, \ldots; \boldsymbol{\theta}) \\
&\propto p(\mathbf{Y}^{s,1} | \mathbf{U}^s; \boldsymbol{\theta}_{E1}) \times p(\mathbf{Y}^{s,2} | \mathbf{U}^s; \boldsymbol{\theta}_{E2}) \times \ldots \times p(\mathbf{U}^s; \boldsymbol{\theta}_A). && (4)
\end{aligned}$$

This yields the expected latent assignments $\langle \mathbf{U}^s \rangle_q$, which serve as estimates of individual-specific parcellations. In the M-step, the parameters $\boldsymbol{\theta}$ are updated by maximizing the ELBO given $\langle \mathbf{U}^s \rangle_q$. The E- and M-steps are iterated until convergence (see section S8).

#### S2 Spatial arrangement model

The spatial arrangement model defines prior distributions $p(\mathbf{U}; \boldsymbol{\theta}_A)$ and $p(\mathbf{U}^s; \boldsymbol{\theta}_A)$ over group-level and individual-specific cortical parcellations $\mathbf{U}$ and $\mathbf{U}^s$. In this work, we introduce mRBM as the spatial arrangement model, which explicitly captures spatial dependencies among neighboring cortical locations and learns the group-level probability that location $i$ belongs to network $k$, denoted $p(\mathbf{u}_i = k)$. Under this model, the posterior assignment probability at location $i$ is given by:

$$p(\mathbf{u}_i = k|\mathbf{y}_i; \boldsymbol{\theta}_A) = \frac{\exp(\ell_{i,k} + \eta_{i,k} + \mathbf{H}_k \mathbf{W}_i \cdot \theta_w)}{\sum_l \exp(\ell_{i,k} + \eta_{i,l} + \mathbf{H}_l \mathbf{W}_i \cdot \theta_w)}, \tag{5}$$

where $\mathbf{W}$, $\mathbf{H}$, and $\theta_w$ are model parameters. For notational simplicity, individual and dataset indices $s$ and $n$ are omitted. The mRBM model is parameterized by a group-level log-probability term $\eta_{i,k}$, which captures the baseline tendency for location $i$ to be assigned to network $k$ across individuals. Eq. 5 can be interpreted as the expected value of the parcellation layer under the mRBM model, approximated via Gibbs sampling in the negative phase (see Eq. 22), averaged across individuals. In the absence of a Gibbs chain (e.g., during initialization), the group-level assignment probabilities reduce to a softmax over $\eta_{i,k}$: $p(\mathbf{u}_i = k) = \exp(\eta_{i,k}) / \sum_j \exp(\eta_{i,j})$, which provides a group-level atlas independent of individual-specific data.

#### S3 Dataset-specific emission models

Features derived from fMRI data $\mathbf{y}_i$ (e.g., resting-state functional connectivity or task-evoked activation profiles) can vary substantially across datasets, participants, and cortical locations. As a result, two vertices within the same functional region may exhibit highly similar measurement profiles but markedly different signal magnitudes. To better capture such directional similarity, we model the data using a mixture of von Mises-Fisher (vMF) distributions (Banerjee et al., 2005; Yeo et al., 2011; Ryali et al., 2013; Schaefer et al., 2018), which has been shown to outperform Gaussian mixture models for both resting-state and task-based fMRI (Røge et al., 2017; Zhi et al., 2025).

The probability density function of an $N$-dimensional ($N \geqslant 2$) vMF distribution for a unit-norm data vector $\mathbf{y}_i$ ($\|\mathbf{y}_i\| = 1$) is:

$$p_N(\mathbf{y}_i|\mathbf{v}, \kappa) = c_N(\kappa) \cdot \exp(\kappa \mathbf{v}^\intercal \mathbf{y}_i), \tag{6}$$

where $\mathbf{v}$ is the mean direction with $\|\mathbf{v}\| = 1$ and $\kappa \geqslant 0$ is the concentration parameter. Larger values of $\kappa$ correspond to tighter concentration around the mean direction. The normalizing constant $c_N(\kappa)$ is given by:

$$c_N(\kappa) = \frac{\kappa^{\frac{N}{2}-1}}{(2\pi)^{\frac{N}{2}} I_{\frac{N}{2}-1}(\kappa)}, \tag{7}$$

where $I_r(\cdot)$ denotes the modified Bessel function of the first kind of order $r$.

In a vMF mixture model with $K$ components, each network $k \in \{1, 2, \cdots, K\}$ is parameterized by $\{\mathbf{v}_k, \kappa_k\}$. Assuming spatial independence of measurement noise, the data log-likelihood for individual $s$ in dataset $n$ at cortical location $i$ is:

$$\ell_{i,k}^{s,n} = \log p(\mathbf{y}_i^{s,n}|\mathbf{u}_i^s(k) = 1; \boldsymbol{\theta}_{En}) = \log c_N(\kappa_k) + \kappa_k \mathbf{v}_k{}^\intercal \mathbf{y}_i^s. \tag{8}$$

We previously investigated three variants of the emission model to accommodate heterogeneity across datasets and sessions (Zhi et al., 2025). In this work, we use the variant that allows different sessions from the same individual to have distinct concentration parameters and models each session with its own emission model (see the illustration of Dataset 2 in Figure 1). Evidence across sessions for the same individual is integrated during the message passing step (see Eq. 4). For identifiability and stability, we assume a shared concentration parameter $\kappa$ across networks within each session. In the M-step, the emission model parameters $\boldsymbol{\theta}_E := \{\mathbf{v}_k, \kappa_k\}$ are updated by maximizing the expected emission log-likelihood $\mathcal{L}_E$ (see section S4).

#### S4 Inference and parameter estimation of the full model

In this section, we describe parameter estimation for the full HBP framework (Algorithm 1). The expected complete-data log-likelihood $\sum_s \langle \log p(\mathbf{Y}^s, \mathbf{U}^s; \boldsymbol{\theta}) \rangle_q$ can be decomposed into an expected emission term $\mathcal{L}_E$ and an

**Algorithm 1:** Model training under the mRBM-HBP framework

**Input:** $K$, fMRI features for individual $s$ and dataset $n$ $\{\mathbf{Y}^{s,n}\}$, initial emission model parameters $\boldsymbol{\theta}_E^{(0)}$, initial arrangement model parameters $\eta_{i,k}^{(0)}$

**Output:** the final estimated parameters $\boldsymbol{\theta}_E^{(t)}$, $\eta_{i,k}^{(t)}$

Initialize: $t = 0$, $t_{\max} = 200$, $\Delta = 1$

**while** $t \leqslant t_{\max}$ **do**

  calculate emission log-likelihoods for each dataset:

  **for** $n = 1$ to $N$ **do**

    emission E-step for each individual $s$ in $\mathbf{Y}^{s,n}$ using Eq.8:

      $\ell_{i,k}^{s,n\,(t)} = \log p(\mathbf{y}_i^{s,n} | \mathbf{u}_i^s(k) = 1; \boldsymbol{\theta}_{En}^{(t)})$

  **end**

  sum emission log-likelihoods across datasets for each individual:

    $\boldsymbol{\ell}_{i,k}^{s\ (t)} = \sum_n \ell_{i,k}^{s,n\,(t)}$

  $\langle \mathbf{U}^s \rangle_+^{(t)}$, $\tilde{\mathbf{U}}^{(t)} \leftarrow$ train mRBM arrangement model using SML (Algorithm 2)

  calculate convergence criterion $\mathcal{J}^{(t)}$ by Eq.29

  **if** $t \geqslant 1$ **and** $\mathcal{J}^{(t)} - \mathcal{J}^{(t-1)} < \Delta$ **then**

    **return** $\langle \mathbf{U}^s \rangle_+^{(t)}$ (individual parcellations), $\tilde{\mathbf{U}}^{(t)}$ (group parcellation)

  **end**

  **for** $n = 1$ to $N$ **do**

    emission M-step by Eq.11 and 12

      $\boldsymbol{\theta}_{En}^{(t+1)} \leftarrow \text{argmax}_{\boldsymbol{\theta}_{En}} \mathcal{L}_{En}^{(t)}(\boldsymbol{\theta}_{En})$

  **end**

  $t \leftarrow t + 1$

**end**

expected arrangement term $\mathcal{L}_A$, where $\langle \cdot \rangle_q$ denotes expectation with respect to the variational distribution $q$. Correspondingly, the model parameters partition as $\boldsymbol{\theta} = \{\boldsymbol{\theta}_E, \boldsymbol{\theta}_A\}$, enabling separate updates within the emission and arrangement components (see section S1). This model structure leads to the following iterative learning procedure.

**Step 1: Emission model E-step**. We represent a given dataset $\mathbf{Y}^n$ as a tensor of size $S \times N \times P$, where $S$ is the number of individuals, $N$ is the number of observations (e.g., task conditions), and $P$ is the number of cortical locations. For each individual $s$, the imaging feature at location $i$, $\mathbf{y}_i^s$, is an $N$-dimensional vector. When repeated measurements of $M$ task conditions are available, we optionally summarize the data using a design matrix $X \in \mathbb{R}^{N \times M}$. To account for multiple partitions (e.g., runs or sessions), we divide the observations into $J$ subsets and construct a combined representation, which is the sum of normalized data across partitions:

$$\tilde{\mathbf{y}}_i^s = \sum_{j=1}^{J} \|(\mathbf{X}_j^\top \mathbf{X}_j)^{-1} \mathbf{X}_j^\top \mathbf{y}_{i,j}^s\|. \tag{9}$$

Alternatively, partitions can be treated as independent observations and normalized separately. Using $\tilde{\mathbf{y}}_i^s$, the expected emission log-likelihood with a $K$-component vMF distribution at iteration $t+1$ is:

$$\mathcal{L}_E{}^{(t+1)} = SPJ \sum_k \log c_M(\kappa_k^{(t)}) + \sum_{s \in \mathbb{S}} \sum_{i=1}^{P} \sum_{k=1}^{K} \langle \mathbf{u}_i^s(k) \rangle_q^{(t)} \kappa_k^{(t)} \mathbf{v}_k^{(t)}{}^\top \tilde{\mathbf{y}}_i^s \tag{10}$$

Note that $\tilde{\mathbf{y}}_i^s$ is $M$-dimensional and not unit-normalized. Accordingly, the normalization constant is computed in $M$ dimensions, denoted as $\log c_M(\kappa_k)$.

**Step 2: Learning the mRBM spatial arrangement model**. The mRBM model does not admit a closed-form posterior. We therefore train the model using stochastic maximum likelihood (SML) with contrastive divergence. This step takes the aggregated emission log-likelihoods from Step 1 as input and returns sufficient statistics (e.g., expected assignments) for subsequent M-step emission model update. Details are provided in section S6 (Algorithm 2).

**Step 3: Emission model M-step**. In the M-step, we update the emission parameters $\boldsymbol{\theta}_E = \{\mathbf{v}_k, \kappa_k\}$ by maximizing $\mathcal{L}_E$. The mean direction $\mathbf{v}_k$ has a closed-form update:

$$\mathbf{v}_k^{(t+1)} = \frac{\tilde{\mathbf{v}}_k}{r_k}, \quad \text{where } \tilde{\mathbf{v}}_k = \sum_s \sum_i \langle \mathbf{u}_i^{s(k)} \rangle_q^{(t)} \cdot \tilde{\mathbf{y}}_i^s, \quad r_k = ||\tilde{\mathbf{v}}_k||. \tag{11}$$

Updating the concentration parameter $\kappa_k$ is more challenging in the high-dimensional setting, as it involves inverting ratios of Bessel functions. Following Banerjee et al. (2005) and Hornik and Grün (2014), we use an approximation under the assumption that the concentration parameter $\kappa$ is shared across networks:

$$\kappa^{(t+1)} \approx \frac{\bar{r}M - \bar{r}^3}{1 - \bar{r}^2}, \tag{12}$$

$$\bar{r} = \frac{\sum_{k=1}^{K} \| \sum_{s=1}^{S} \sum_{i=1}^{P} \langle \mathbf{u}_i^{s(k)} \rangle_q^{(t)} \cdot \tilde{\mathbf{y}}_i^s \|}{\sum_{s=1}^{S} \sum_{i=1}^{P} J_i^s}. \tag{13}$$

The above three steps are iterated until convergence.

#### S5 Obtaining group- and individual-level parcellations

Group-level parcellations can be estimated using resting-state functional connectomes, task-evoked activation profiles, or fused datasets. In practice, we observed minimal differences between the spatially independent and mRBM-based arrangement models at the group level. Therefore, for computational efficiency and simplicity, we use the spatially independent model (Zhi et al., 2025), which is equivalent to the mRBM-HBP model with $\theta_w = 0$. Under this formulation, the group atlas is represented by network assignment probabilities at each cortical location,

$$p(\mathbf{u}_i) = \text{softmax}(\boldsymbol{\eta}_i),$$

which are assembled into the matrix $\mathbf{U}$.

To account for imbalances in the amount of data across datasets and modalities, an internal weighting scheme was applied during training. Specifically, when aggregating emission log-likelihoods across datasets in the collaborative learning step (Algorithm 1, line 9), we weighted $\ell_{i,k}^{s,n(t)}$ by the inverse of the dimensionality of the data. This ensures balanced contributions from each dataset.

Individual-level parcellations are inferred by combining the learned group-level prior with individual-specific data. After training the full model, a new dataset can be incorporated to compute the expected value of $\mathbf{U}^s$, yielding a probabilistic parcellation for each individual (see section S8). In practice, we first fit a new dataset-specific emission model to the individual-level data while keeping the parameters of the spatial arrangement model fixed. Following training via stochastic maximum likelihood (SML; see section S6), individual-level parcellations are obtained through a single E-step that integrates the individual data likelihood with the group prior. The posterior distribution can be expressed as:

$$p(\mathbf{U}^s | \mathbf{Y}^s; \boldsymbol{\theta}_A, \boldsymbol{\theta}_E) = \text{softmax}(\boldsymbol{\ell}^s + \boldsymbol{\eta} + \mathbf{H}^s \mathbf{W} \cdot \theta_w). \tag{14}$$

For comparison in simulation analyses (section S10), we also consider a data-only parcellation that is not informed by the group prior:

$$p(\mathbf{U}^s|\mathbf{Y}^s, \boldsymbol{\theta}_E) = \text{softmax}(\boldsymbol{\ell}^s). \tag{15}$$

#### S6 Training the mRBM model using stochastic maximum likelihood

The architecture of the mRBM model enables parameter estimation within a variational framework using stochastic maximum likelihood (Salakhutdinov and Hinton, 2012). Model parameters are updated via contrastive divergence (CD) (Hinton et al., 2006), which provides an efficient approximation to the gradient of the unnormalized expected log-likelihood of the spatial arrangement model (Eq. 17). In the positive phase (section S6.1), the data distribution is approximated using a mean-field factorization:

$$q(\mathbf{U}^s, \mathbf{H}^s|\mathbf{Y}^s) \approx \prod_i p(\mathbf{u}_i^s|\mathbf{Y}^s) \prod_j p(\mathbf{h}_j^s|\mathbf{U}^s), \tag{16}$$

where $\mathbf{U}^s$ and $\mathbf{H}^s$ denote the parcellation and hidden layers, respectively. In the negative phase, samples are drawn by alternating Gibbs updates from the conditional distributions $p(\mathbf{Y}^s|\mathbf{U}^s)$, $p(\mathbf{H}^s|\mathbf{U}^s)$, and $p(\mathbf{U}^s|\mathbf{H}^s, \mathbf{Y}^s)$ (section S6.2). The model parameters $\boldsymbol{\theta}_A := \{\boldsymbol{\eta}, \theta_w\}$ are subsequently updated in the parameter update step (section S6.3).

To integrate mRBM training into the HBP framework, we use SML with mini-batch learning (Algorithm 2), corresponding to line 10 in Algorithm 1. This design preserves the standard E- and M-steps of the emission model while incorporating mRBM-based updates for the arrangement model. For notational simplicity, we omit the dataset index $n$ in the algorithm. The softmax$(\cdot)$ function serves as the activation function for each multinomial node and is applied along the node dimension for matrix inputs.

The SML procedure takes as input the data log-likelihoods of all individuals in $\mathbb{S}$. After convergence, it yields the approximate posterior expectations $\langle \mathbf{u}_i^s(k)\rangle_q$ (denoted $\langle \mathbf{U}^s\rangle_+$) for each individual. The unnormalized expected arrangement log-likelihood of the mRBM model is:

$$\begin{aligned}\widetilde{\mathcal{L}}_A &= \sum_{s\in\mathbb{S}} \langle \log \tilde{p}(\mathbf{U}^s; \boldsymbol{\theta}_A)\rangle_q \\ &= \sum_{s\in\mathbb{S}} \sum_{i,k} \langle \mathbf{u}_i^s(k)\rangle_q \cdot \eta_{i,k} + \sum_{s\in\mathbb{S}} \sum_{i,j,k} \theta_w \cdot w_{i,j} \langle \mathbf{u}_i^s(k)\rangle_q \langle \mathbf{h}_j^s(k)\rangle_q. \end{aligned} \tag{17}$$

Parameter updates follow the gradient of $\widetilde{\mathcal{L}}_A$ with respect to $\boldsymbol{\theta}_A$. In practice, we approximate these gradients using persistent contrastive divergence (PCD; or PCD-$k$ with $k$ Gibbs steps per update) (Tieleman, 2008), decomposing learning into positive (section S6.1) and negative (section S6.2) phases. The resulting parameter updates are then applied as described in section S6.3.

##### S6.1 Positive phase: expectation given the data

In the positive phase, we approximate the data-dependent posterior distribution using a mean-field factorization (Eq. 16). Expectations under this distribution are denoted by $\langle\cdot\rangle_+$.

At iteration $t$, the posterior expectation of the parcellation assignment for cortical location $i$ in individual $s$ is:

$$\begin{aligned}\langle \mathbf{u}_i^s(k)\rangle_+^{(m)} &= p(\mathbf{u}_i^s = k|\mathbf{y}_i^s; \boldsymbol{\theta}_A^{(t)}, \boldsymbol{\theta}_E^{(t)}) \\ &= \frac{\exp(\langle \log p(\mathbf{y}_i^s|\mathbf{u}_i^s = k; \boldsymbol{\theta}_E^{(t)})\rangle_+ + \eta_{i,k}{}^{(t)} + \langle \mathbf{H}_{k,\cdot}^s\rangle_+^{(m)} \mathbf{W}_{\cdot,i} \cdot \theta_w^{(t)})}{\sum_l \exp(\langle \log p(\mathbf{y}_i^s|\mathbf{u}_i^s = l; \boldsymbol{\theta}_E^{(t)})\rangle_+ + \eta_{i,l}{}^{(t)} + \langle \mathbf{H}_{l,\cdot}^s\rangle_+^{(m)} \mathbf{W}_{\cdot,i} \cdot \theta_w^{(t)})},\end{aligned} \tag{18}$$

where $m$ denotes the number of mean-field iterations. The hidden layer expectations are updated as:

$$\langle \mathbf{h}_j^s(k)\rangle_+^{(m+1)} = p(\mathbf{h}_j^s = k|\langle \mathbf{U}^s\rangle_+, \boldsymbol{\theta}_A^{(t)}) = \frac{\exp(\langle \mathbf{U}_{k,\cdot}^s\rangle_+^{(m)} \mathbf{W}_{j,\cdot}^\intercal \cdot \theta_w^{(t)})}{\sum_l \exp(\langle \mathbf{U}_{l,\cdot}^s\rangle_+^{(m)} \mathbf{W}_{j,\cdot}^\intercal \cdot \theta_w^{(t)})}. \tag{19}$$

**Algorithm 2:** The stochastic maximum likelihood (SML) algorithm for the mRBM model.

**Input:** a collection of individual-specific data likelihoods $\mathbb{L} = \{\boldsymbol{\ell}_{i,k}^{1\ (t)}, ..., \boldsymbol{\ell}_{i,k}^{s\ (t)}\}, s \in \mathbb{S}$, and corresponding persistent Gibbs chains for the negative phase $\tilde{\mathbf{U}}^{s^{(0)}}$

**Output:** Posterior expectations $\langle \mathbf{U}^s \rangle_+^{(t)}$, i.e., $\langle \mathbf{u}_i^s(k) \rangle_q^{(t)}$ for all individuals, and updated Gibbs chains $\tilde{\mathbf{U}}^{s^{(n)}}$

**Set** learning rate $\alpha$ to a small positive value; mini-batch size $B$; $m = 5$ mean-field iterations in the positive phase, $n = 10$ Gibbs sampling steps in the negative phase

**Initialize** counter$= 1$; two empty $S \times K \times P$ tensors, $\tilde{\mathbf{Y}}$ and $\tilde{\mathbf{H}}$, for samples in the negative phase; set $\langle \mathbf{H} \rangle_+^{(0)} = 0$ for the positive phase; if persistent chains are not provided, initialize $\tilde{\mathbf{U}}^{(0)}$ by sampling from softmax$(\boldsymbol{\eta}^{(t)})$

**while** counter $\leqslant \lceil S/B \rceil$ **do**

  Randomly sample a mini-batch of $\mathbb{B}$ from $\mathbb{L}$ without replicates. Remove $\mathbb{B}$ from $\mathbb{L}$.

  (1) Positive phase: perform $m$ iterations of mean-field approximation using Eq.18 and 19:

  **for** $a = 0$ to $m - 1$ **do**

    $\forall s \in \mathbb{B},\ \langle \mathbf{U}^s \rangle_+^{(a)} = \text{softmax}(\boldsymbol{\ell}^{s\,(t)} + \boldsymbol{\eta}^{(t)} + \langle \mathbf{H}^s \rangle_+^{(a)} \mathbf{W} \cdot \theta_w^{(t)})$

    $\forall s \in \mathbb{B},\ \langle \mathbf{H}^s \rangle_+^{(a+1)} = \text{softmax}(\langle \mathbf{U}^s \rangle_+^{(a)} \mathbf{W}^\intercal \cdot \theta_w^{(t)})$

  **end**

  $\langle \mathbf{U}^s \rangle_+^{(t)} \leftarrow \langle \mathbf{U}^s \rangle_+^{(m-1)} \quad (s \in \mathbb{B}), \quad \langle \mathbf{H}^s \rangle_+^{(t)} \leftarrow \langle \mathbf{H}^s \rangle_+^{(m)} \quad (s \in \mathbb{B})$

  (2) Negative phase: $n$ Gibbs mixing steps using Eq.22 and 21:

  **for** $b = 0$ to $n - 1$ **do**

    $\forall s \in \mathbb{B},\ \tilde{\mathbf{Y}}^{s^{(b)}}$ sampled from $p(\mathbf{Y}^s | \tilde{\mathbf{U}}^{s^{(b)}}; \boldsymbol{\theta}_E^{(t)})$

    $\forall s \in \mathbb{B},\ \tilde{\mathbf{H}}^{s^{(b)}}$ sampled from softmax$(\tilde{\mathbf{U}}^{s^{(b)}} \mathbf{W}^\intercal \cdot \theta_w^{(t)})$

    $\forall s \in \mathbb{B},\ \tilde{\mathbf{U}}^{s^{(b+1)}}$ sampled from softmax$(\tilde{\boldsymbol{\ell}}^{s^{(b)}} + \boldsymbol{\eta}^{(t)} + \tilde{\mathbf{H}}^{s^{(b)}} \mathbf{W} \cdot \theta_w^{(t)})$

  **end**

  $\langle \mathbf{U}^s \rangle_-^{(t)} \leftarrow \tilde{\mathbf{U}}^{s^{(n)}} \quad (s \in \mathbb{B}), \quad \langle \mathbf{H}^s \rangle_-^{(t)} \leftarrow \tilde{\mathbf{H}}^{s^{(n-1)}} \quad (s \in \mathbb{B})$

  $\tilde{\mathbf{U}}^{s^{(0)}} \leftarrow \langle \mathbf{U}^s \rangle_-^{(t)} \quad (s \in \mathbb{B})$ // Reset $\tilde{\mathbf{U}}^{s^{(0)}}$ for persistent-CD in the next iteration

  (3) Parameters update using Eq.26 and 27:

  $$\theta_w^{(t)} = \theta_w^{(t)} + \alpha \cdot \frac{1}{B} \mathbf{W} \sum_{s \in \mathbb{B}} (\langle \mathbf{H}^s \rangle_+^{(t)\intercal} \langle \mathbf{U}^s \rangle_+^{(t)} - \langle \mathbf{H}^s \rangle_-^{(t)\intercal} \langle \mathbf{U}^s \rangle_-^{(t)})$$

  $$\boldsymbol{\eta}^{(t+1)} = \boldsymbol{\eta}^{(t)} + \alpha \cdot \frac{1}{B} \sum_{s \in \mathbb{B}} (\langle \mathbf{U}^s \rangle_+^{(t)} - \langle \mathbf{U}^s \rangle_-^{(t)})$$

  counter $\leftarrow$ counter $+1$

**end**

**return** $\langle \mathbf{U} \rangle_+^{(t)}, \tilde{\mathbf{U}}^{(0)}$

We initialize $\langle \mathbf{H}^s \rangle_+^{(0)} = 0$ and alternate updates of $\langle \mathbf{U}^s \rangle_+$ and $\langle \mathbf{H}^s \rangle_+$ for $m$ iterations. The final estimates are recorded as: $\langle \mathbf{U}^s \rangle_+^{(t)} = \langle \mathbf{U}^s \rangle_+^{(m)}$ and $\langle \mathbf{H}^s \rangle_+^{(t)} = \langle \mathbf{H}^s \rangle_+^{(m+1)}$.

##### S6.2 Negative phase: expectation given the model

In the negative phase, we approximate expectations under the model distribution using $n$ steps of Gibbs sampling. Expectations under this distribution are denoted by $\langle \cdot \rangle_-$. At iteration $t$, the mixing of samples $\tilde{\mathbf{U}}^s$ and $\tilde{\mathbf{H}}^s$ for each individual $s$ is initialized using a persistent multinomial Markov chain (Tieleman, 2008) over the parcellation layer. If no persistent chain is available, the initial state is sampled as $\tilde{\mathbf{U}}^{s^{(0)}} \sim \text{softmax}(\boldsymbol{\eta}^{(t)})$.

At the $n^{th}$ Gibbs iteration, the sampling procedure consists of the following three steps.

(1) Data sampling. Synthetic observations are drawn from the emission model:

$$\tilde{\mathbf{Y}}^{s^{(n)}} \sim p(\mathbf{Y}^s | \tilde{\mathbf{U}}^{s^{(n)}}; \boldsymbol{\theta}_E^{(t)}), \tag{20}$$

and the corresponding data log-likelihood $\tilde{\boldsymbol{\ell}}^{s(n)}$ is computed using the emission model (Eq. 8).

(2) Hidden layer update. Due to the bipartite structure of mRBM, the hidden layer can be sampled conditional on the parcellation layer:

$$\tilde{\mathbf{H}}^{s(n)} \sim p(\mathbf{H}^s|\tilde{\mathbf{U}}^{s(n)}; \boldsymbol{\theta}_A^{(t)}) = \text{softmax}(\tilde{\mathbf{U}}^{s(n)}\mathbf{W}^\intercal \cdot \theta_w^{(t)}), \tag{21}$$

where the resulting $K \times J$ matrix defines a collection of multinomial distributions, one for each hidden node. Each component $\tilde{\mathbf{h}}_j^{s(n)}$ is sampled independently from its corresponding distribution and assembled into $\tilde{\mathbf{H}}^{s(n)}$.

(3) Parcellation layer update. Lastly, the parcellation layer is updated conditioned on both the sampled data and the hidden states:

$$\tilde{\mathbf{U}}^{s(n)} \sim p(\mathbf{U}^s|\tilde{\mathbf{Y}}^{s(n)}, \tilde{\mathbf{H}}^{s(n)}; \boldsymbol{\theta}_A^{(t)}) = \text{softmax}(\tilde{\boldsymbol{\ell}}^{s(n)} + \boldsymbol{\eta}^{(t)} + \tilde{\mathbf{H}}^{s(n)}\mathbf{W} \cdot \theta_w^{(t)}). \tag{22}$$

The resulting $K \times P$ matrix defines multinomial distributions for each parcellation node, from which $\tilde{\mathbf{u}}_i^{s(n)}$ are sampled independently.

In the final Gibbs iteration, we use the probabilities directly, rather than sampling multinomial realizations, for both $\tilde{\mathbf{U}}^{s(n+1)}$ and $\tilde{\mathbf{H}}^{s(n)}$ to reduce sampling variance (Hinton, 2012). These are then recorded as sufficient statistics for subsequent gradient computation:

$$\langle \mathbf{U}^s \rangle_-^{(t)} = \tilde{\mathbf{U}}^{s(n+1)}, \quad \langle \mathbf{H}^s \rangle_-^{(t)} = \tilde{\mathbf{H}}^{s(n)}. \tag{23}$$

The updated $\tilde{\mathbf{U}}^{s(n+1)}$ is retained as the initial state for the next iteration $(t+1)$ of persistent contrastive divergence (Tieleman, 2008).

##### S6.3 Update phase: parameter estimation using gradients

The mRBM model parameters are updated by following the gradients of the unnormalized expected arrangement log-likelihood in Eq. 17. Given the positive- and negative-phase expectations of the latent and hidden variables, the gradients with respect to $\boldsymbol{\theta}_A := \{\eta_{i,k}, \theta_w\}$ at iteration $t$ are:

$$\nabla \theta_w^{(t)} = \frac{1}{S}\mathbf{W}\sum_s (\langle \mathbf{H}^s \rangle_+^{(t)\intercal} \langle \mathbf{U}^s \rangle_+^{(t)} - \langle \mathbf{H}^s \rangle_-^{(t)\intercal} \langle \mathbf{U}^s \rangle_-^{(t)}), \tag{24}$$

$$\nabla \boldsymbol{\eta}^{(t)} = \frac{1}{S}\sum_s (\langle \mathbf{U}^s \rangle_+^{(t)} - \langle \mathbf{U}^s \rangle_-^{(t)}). \tag{25}$$

The parameters are then updated via gradient ascent:

$$\theta_w^{(t+1)} = \theta_w^{(t)} + \alpha \cdot \nabla \theta_w^{(t)}, \tag{26}$$

$$\boldsymbol{\eta}^{(t+1)} = \boldsymbol{\eta}^{(t)} + \alpha \cdot \nabla \boldsymbol{\eta}^{(t)}, \tag{27}$$

where $\alpha$ is the learning rate, which is set to a small positive value.

#### S7 Practical guidance for training the mRBM model

As the mRBM model is derived from the restricted Boltzmann machine (RBM) family, it inherits limitations associated with training via contrastive divergence (Hinton, 2002). Consequently, careful implementation choices and practical considerations are required to ensure stable and efficient model training. In this section, we describe the key algorithmic settings and techniques used to improve training performance.

Our primary training framework is stochastic maximum likelihood (SML), also known as persistent contrastive divergence (PCD) (Tieleman, 2008), combined with mini-batch learning. Compared to standard contrastive divergence (CD), SML maintains and updates persistent Markov chains across iterations rather than reinitializing them at each step. This allows for more efficient exploration of the model distribution and typically leads to faster and more stable convergence, particularly for complex models (Tieleman, 2008). The use of mini-batches further introduces stochasticity by sampling different subsets of training data at each iteration, which helps the optimization escape shallow local minima. In addition, mini-batch training reduces memory usage and computational cost, enabling efficient GPU-accelerated matrix operations for large-scale imaging data.

We also incorporated several standard training heuristics (Hinton, 2012), including careful tuning of batch size, learning rate, number of training epochs, momentum, and the number of Gibbs sampling steps. In practice, these hyperparameters are interdependent. Due to GPU memory constraints, we first determined the largest mini-batch size $B$ by selecting the largest divisor of the total number of individuals that could be accommodated in memory; if necessary, smaller divisors were considered iteratively. At the beginning of each epoch, $B$ individuals were randomly sampled, and their corresponding data likelihoods were used for computation. After processing each mini-batch, individual-specific sufficient statistics were mapped to their original indices. Sampling was performed without replacement within each epoch, ensuring that all individuals were used exactly once per epoch.

The total number of training epochs was fixed at $t = 200$, which served as the stopping criterion due to the lack of a reliable convergence diagnostic for the model. The base learning rate was set to a relatively small value $\alpha = 0.01$ to prevent overshooting the optimum, at the cost of potentially slower convergence. To mitigate this, we incorporated momentum (Sutskever et al., 2013), which accelerates learning in directions of consistent gradient descent and is particularly effective in navigating objective functions with narrow, elongated valleys. Specifically, Eq. 26 and 27 were replaced with the following parameter updates:

$$\boldsymbol{\theta}_A^{(t+1)} = \boldsymbol{\theta}_A^{(t)} + \boldsymbol{v}^{(t+1)}, \quad \boldsymbol{v}^{(t+1)} = \gamma \boldsymbol{v}^{(t)} - \alpha \nabla \boldsymbol{\theta}_A^{(t)}, \tag{28}$$

where $\boldsymbol{v}^{(t)}$ denotes the velocity term and $\gamma$ is the momentum coefficient controlling the contribution of past gradients. In this work, we set $\gamma = 0.9$.

Finally, we set the number of mean-field iterations in the positive phase to $m = 5$ (section S6.1) and the number of Gibbs sampling steps in the negative phase to $n = 10$ (section S6.2). These values were chosen to provide sufficient burn-in for the Markov chains to approach equilibrium while maintaining computational efficiency.

#### S8 Monitoring learning progress, convergence, and model fine-tuning

To monitor the learning progress of the full model, we constructed a pseudo-objective function $\mathcal{J}$ during training. Since the mRBM model does not admit a closed-form posterior, $\mathcal{J}$ combines the expected emission log-likelihood $\mathcal{L}_E$ with a negative cross-entropy term $\mathcal{H}(\cdot)$ between the positive- and negative-phase expectations, $\langle \mathbf{U} \rangle_+^{(t)}$ and $\langle \mathbf{U} \rangle_-^{(t)}$. For a single dataset or session, $\mathcal{J}$ is defined as:

$$\mathcal{J}^{(t)} = -\sum_{s \in \mathbb{S}} \mathcal{H}(\langle \mathbf{U}^s \rangle_+^{(t)}, \langle \mathbf{U}^s \rangle_-^{(t)}) + \sum_{s \in \mathbb{S}} \langle \log p(\mathbf{Y}^s | \mathbf{U}^{s(t)}; \boldsymbol{\theta}_E{}^{(t)}) \rangle_q, \tag{29}$$

where the first term penalizes discrepancies between the reconstructed latent variables $\langle \mathbf{U}^s \rangle_-$ and the data-driven expectations $\langle \mathbf{U}^s \rangle_+$. When the two match exactly, this term vanishes and $\mathcal{J}$ reduces to $\mathcal{L}_E$. In practice, a monotonic increase in $\mathcal{J}^{(t)}$ provides an empirical indicator that the learning process is progressing in a favorable direction.

Training the mRBM model may exhibit convergence instability, potentially yielding multiple plausible parameter configurations (Zhi, 2023). To mitigate the propagation of such variability to downstream individual-level inference, we fixed the group-level probability parameter $\boldsymbol{\eta}$ to the estimate obtained from the spatially independent model. The temperature parameter $\theta_w$ was treated as a tunable hyperparameter within a positive range (from 0.1 to 15). This temperature parameter controls the degree of spatial smoothness in the inferred parcellations, with larger values producing smoother functional boundaries.

For individual-level inference, we further introduced a scaling parameter $\lambda$ to control the relative contribution of the individual data versus the group prior. Specifically, the expected complete log-likelihood (Eq. 3) for individual $s$ was modified as:

$$\mathcal{L}^s = \langle \log p(\mathbf{Y}^s, \mathbf{U}^s; \boldsymbol{\theta}) \rangle_q = \langle \log p(\mathbf{Y}^s | \mathbf{U}^s; \boldsymbol{\theta}_E) \rangle_q + \lambda \langle \log p(\mathbf{U}^s; \boldsymbol{\theta}_A) \rangle_q \triangleq \mathcal{L}_E^s + \lambda \mathcal{L}_A^s, \tag{30}$$

where $\lambda$ scales the influence of the group prior relative to the individual data. This modification improves robustness, as individual-level fMRI data are often limited in duration and vary in signal-to-noise ratio across datasets and acquisition protocols. Larger values of $\lambda$ place greater weight on the group prior, whereas smaller values emphasize individual-specific data.

To determine optimal hyperparameters $\theta_w$ and $\lambda$ for the HCP-YA dataset, we performed grid search using a validation set and evaluated parcellation quality on independent task activation maps using DCBC and task inhomogeneity metrics. The selected parameters were then applied to infer parcellations for test individuals. For the HPN dataset, which includes only 15 participants, hyperparameter tuning was conducted directly on the available individuals using independent runs rather than using a separate validation set. In this case, only $\lambda$ was tuned, as the data quality was relatively high. We fixed $\theta_w = 0$ to minimize imposed spatial smoothing and preserve individual-specific functional boundary information.

#### S9 Correction for left-right hemisphere symmetry in the fs-LR32k template

All parcellation analyses in this study were performed using the HCP fs-LR32k template (Van Essen et al., 2012), which contains 32,492 vertices per hemisphere. However, in the standard template, the medial wall is not perfectly symmetric, leading to a mismatch in the number of cortical grayordinates between the left (29,696) and right (29,716) hemispheres. To enforce strict hemispheric symmetry and enable vertex-wise correspondence for downstream analyses (e.g., lateralization studies), we applied a manual correction. Specifically, we redefined the medial wall as the intersection of medial wall vertices across the left and right hemispheres, thereby maximizing the preservation of cortical data. After this adjustment, the template contained 29,759 spatially corresponding cortical vertices in each hemisphere.

#### S10 Validating mRBM-HBP via simulation

##### S10.1 Synthetic data generation for simulation

To validate the proposed spatial arrangement model, we conducted a series of simulations using synthetic datasets. The data generation procedure largely followed Zhi et al. (2025), with several modifications tailored to the current work. Specifically, we used a four-neighbor Markov Random Field (MRF) defined on a $50 \times 50$ grid to generate ground-truth individual-level parcellations. Each vertex on the grid represents a location assigned to one of $K$ parcels. Throughout this study, we used $K = 5$.

We first constructed a group-level probability map using five centroids $\mu_k$, $k = 1, 2, \cdots, 5$, located at four corners and the center of the MRF grid (Supplementary Figure 1a). The group-level log-probability (bias term) for parcel $k$ at location $x_i$ is defined as:

$$\eta_{i,k} = -\frac{|x_i - \mu_k|_2^2}{2\sigma_\mu^2}, \tag{31}$$

where $\sigma_\mu^2$ controls the spatial smoothness of the group map (Supplementary Figure 1a). The group-level probability $p(\mathbf{u}_i)$ were then obtained by applying the `softmax` function to $\boldsymbol{\eta}_i$, yielding a multinomial distribution over parcels at each location.

Initial individual parcellation maps $\mathbf{U}^s$ were generated by sampling parcel assignments independently at each location according to $p(\mathbf{u}_i)$. These initial maps were subsequently refined using a modified mRBM model via layer-wise Gibbs sampling to introduce spatial dependencies. The mRBM model consisted of a parcellation layer and a hidden layer, each containing $P$ nodes (the number of grid vertices), connected by symmetric pairwise weights $w_{i,j} = w_{j,i}$.

The weights encoded local spatial structure, with $w_{i,j} = 1$ if vertices $i$ and $j$ are neighbors and $w_{i,j} = 0$ otherwise. The spatial coupling strength was controlled by a temperature parameter $\theta_w$, such that larger values encourage neighboring vertices to share the same parcel assignment, thereby enforcing local smoothness (Supplementary Figure 1c). For simplicity, bias terms in both the parcellation and hidden layers were omitted in the synthetic data generation process. After $m$ burn-in iterations of alternating Gibbs sampling between parcellation and hidden layers (Supplementary Figure 1c), stable individual parcellation maps were obtained (Supplementary Figure 1d).

Given the individual parcellations, observed synthetic data $\mathbf{Y}^s$ were generated for each individual. Specifically, we used an $N$-dimensional Gaussian mixture model (GMM) with component means $\mathbf{v}_k$ normalized to unit length ($\|\mathbf{v}_k\| = 1$). For each vertex $i$ assigned to parcel $k$, the observed data were generated as:

$$\mathbf{y}_i = \mathbf{v}_k + \boldsymbol{\epsilon}, \tag{32}$$

where $\boldsymbol{\epsilon}$ was Gaussian noise with variance $\sigma_k^2\mathbf{I}$. This process yielded a synthetic dataset with $N$ observations per vertex across $P$ locations and $S$ individuals.

For the simulation results presented in section S10.2, we further applied spatial smoothing to the synthetic data using a $5 \times 5$ Gaussian kernel with varying standard deviation $\sigma$, convolving the data over the MRF grid. The temperature parameter was set to $\theta_w = 1.2$. The group-level bias terms $\eta_{i,k}$ were generated with $\sigma_\mu^2 = 240$. The number of parcels in both the ground-truth and fitted model was fixed at $K = 5$. We generated $S = 30$ individual parcellation maps $\mathbf{U}^s$ using $m = 10$ burn-in iterations with local connectivity weights $\theta_w w_{i,j}$. These maps were then used to generate synthetic training datasets $\{\mathbf{Y}^s\}$ with $N = 10$ observations, as well as independent test sets $\{\mathbf{Y}^s_{\text{test}}\}$ with the same dimensionality.

##### S10.2 The mRBM model effectively captures spatial smoothness

We evaluated the ability of the mRBM model to capture spatial dependencies among cortical locations, in comparison to a spatially independent model. To this end, we conducted simulations using synthetic training datasets generated from the same set of individuals. Model performance was assessed by comparing reconstructed individual parcellations on an independent test set. Evaluation metrics included the mean absolute error between the ground-truth and reconstructed parcellations $\bar{\epsilon}_{|\mathbf{U}-\hat{\mathbf{U}}|}$ (section S10.3), distance-controlled boundary coefficient (DCBC), and the mean adjusted expected cosine error $\bar{\epsilon}_{\text{Acosine}}$ (section S10.4). Each simulation was repeated 100 times using the data generation process described in section S10.1.

The simulations first replicated prior findings (Zhi et al., 2025; Nettekoven et al., 2024) that incorporating group-level information substantially improves individual parcellation estimates relative to using individual data alone. Even under the spatially independent model (Supplementary Figure 2a; *Independent + unsmoothed data*), the inclusion of group priors led to markedly improved reconstruction of the true individual maps compared with the data-only baseline (Supplementary Figure 2a; *Data only*), both qualitatively (Supplementary Figure 2a) and quantitatively (Supplementary Figure 2b). Specifically, the independent model achieved significantly lower mean absolute error ($t_{99} = -1734.16, p = 1.04 \times 10^{-223}$).

Importantly, explicitly modeling spatial dependencies using mRBM yielded further and consistent improvements over the spatially independent model. Visually, comparing with the independent model (Supplementary Figure 2a; *Independent + unsmoothed data*), the mRBM–based parcellations more closely matched the ground-truth maps (Supplementary Figure 2a; *True*) and exhibited sharper parcel boundaries with reduced uncertainty (Supplementary Figure 2a; *mRBM + unsmoothed data*). Quantitatively, mRBM achieved significantly lower mean absolute error than the independent model (Supplementary Figure 2b; $t_{99} = -316.11, p = 1.52 \times 10^{-150}$). This advantage was consistently observed across additional metrics, including mean adjusted expected cosine error (Supplementary Figure 2c; $t_{99} = -298.38, p = 4.59 \times 10^{-148}$) and DCBC (Supplementary Figure 2d; $t_{99} = 220.50, p = 4.44 \times 10^{-135}$).

To assess whether spatial smoothing of the data could compensate for the lack of explicit spatial modeling, we trained the independent model on datasets smoothed at varying levels. Mild smoothing improved performance relative to unsmoothed data, with optimal performance observed at approximately $\sigma = 0.5$ (Supplementary Figure 2b-d). However, even at this optimal smoothing level, the spatially independent model remained significantly inferior to the mRBM model trained on unsmoothed data across all evaluation metrics (Supplementary Figure 2b-d; $\bar{\epsilon}_{|\mathbf{U}-\hat{\mathbf{U}}|}$: $t_{99} = -163.64, p = 2.71 \times 10^{-122}$; $\bar{\epsilon}_{\text{Acosine}}$: $t_{99} = -171.72, p = 2.34 \times 10^{-124}$; DCBC: $t_{99} = -96.92, p = 6.42 \times 10^{-100}$). Moreover, increasing the level of smoothing beyond this optimum led to progressive decrease in performance for the independent model. At higher smoothing levels (e.g., $\sigma = 1.5$), parcellations were substantially degraded and even underperformed compared with those derived from unsmoothed data (Supplementary Figure 2b-d; $\bar{\epsilon}_{|\mathbf{U}-\hat{\mathbf{U}}|}$: $t_{99} = 31.58, p = 1.69 \times 10^{-53}$; $\bar{\epsilon}_{Acosine}$: $t_{99} = 62.53, p = 2.22 \times 10^{-81}$; DCBC: $t_{99} = -64.93, p = 5.80 \times 10^{-83}$).

Overall, these results demonstrate the systematic advantage of the mRBM model over the independent arrangement model in capturing intrinsic spatial dependencies between neighboring locations. Although moderate spatial smoothing can partially mitigate the limitations of the independent model, it cannot substitute for explicit modeling of spatial structure. The mRBM model consistently produced more accurate and spatially contiguous individual parcellations, even when trained on unsmoothed data, highlighting the importance of spatially informed generative models for delineating functional brain organization.

##### S10.3 Mean absolute error

Given access to ground-truth parcel assignments in the simulations, we evaluated the accuracy of reconstruction using the mean absolute error between the true parcellations $\mathbf{U}$ and the estimated probabilistic parcellations $\hat{\mathbf{U}}$. Traditionally, absolute error is computed based on hard parcellations. In this work, we extend this metric to the probabilistic setting by computing the mean expected reconstruction error under the posterior distribution of the estimated parcellations across all cortical locations. Specifically, we define:

$$\langle \bar{\epsilon}_{|\mathbf{U}-\hat{\mathbf{U}}|} \rangle_q = \frac{1}{PS} \sum_{s=1}^{S} \sum_{i=1}^{P} |\mathbf{u}_i^s - \langle \mathbf{u}_i^s \rangle_q|, \tag{33}$$

where $\mathbf{u}_i^s$ denotes the true parcel assignment at location $i$ for individual $s$, represented as a one-hot encoded vector, and $\langle \mathbf{u}_i^s \rangle_q$ denotes the expected parcel assignment under the posterior distribution $q$. Both quantities are multinomial vectors. In practice, this formulation is invariant to label permutation. Because $\langle \mathbf{u}_i^s \rangle_q$ is defined in the same label space as the ground-truth $\mathbf{u}_i^s$, no additional alignment or permutation matching is required, and exhaustive search over label permutations is unnecessary.

##### S10.4 Mean adjusted expected cosine error

To evaluate the predictive power of individual probabilistic parcellations in the simulation, we designed an additional metric, the mean adjusted expected cosine error, $\langle \bar{\epsilon}_{\text{Acosine}} \rangle_q$, defined as:

$$\langle \bar{\epsilon}_{\text{Acosine}} \rangle_q = \frac{1}{\sum_{i=1}^{P} \|\mathbf{y}_i\|^2} \sum_{i=1} \sum_{k=1} \langle \mathbf{u}_i(k) \rangle_q (\|\mathbf{y}_i\|^2 - \mathbf{v}_k{}^T \mathbf{y}_i \|\mathbf{y}_i\|), \tag{34}$$

where $\langle \mathbf{u}_i(k) \rangle_q$ denotes the probabilistic parcellation, i.e., the posterior probability that location $i$ belongs to parcel $k$. The vector $\mathbf{y}_i$ represents the observed profile at location $i$ from the test set $\mathbf{Y}_{\text{test}}$, and $\mathbf{v}_k$ denotes the estimated mean functional response for parcel $k$. A full derivation of this metric, along with its equivalence to computing $1 - R^2$, is provided in section S11.

Conceptually, given the test dataset $\mathbf{Y}_{\text{test}}$, the mean adjusted expected cosine error evaluates how well the probabilistic parcellation can predict unseen data. Specifically, a predicted response $\hat{\mathbf{Y}}_{\text{test}}$ is constructed by combining the probabilistic parcellation $\langle \mathbf{u}_i(k) \rangle_q$ with the estimated parcel-level responses $\mathbf{v}_k$ (Eq. 34). The mean adjusted expected cosine error then quantifies the discrepancy between $\hat{\mathbf{Y}}_{\text{test}}$ and the observed data $\mathbf{Y}_{\text{test}}$ via a cosine-based measure. This formulation provides an interpretable measure of how well each region in the estimated individual parcellation captures observed patterns in the test data. Lower values of $\langle \bar{\epsilon}_{\text{Acosine}} \rangle_q$ indicate better alignment between predicted and observed responses, reflecting more accurate parcellation and more informative parcel-level representations.

#### S11 Mathematical details of the mean adjusted expected cosine error

##### S11.1 Derivation of the mean adjusted expected cosine error

Here we provide the full derivation of the mean adjusted expected cosine error. Because the data are generated or modeled using directional statistical models (e.g., vMF distributions or normalized Gaussian mixed models), cosine-based metrics provide a natural measure of similarity between reconstructed data $\hat{\mathbf{y}}_i$ and observed data $\mathbf{y}_i$. A simple baseline approach is to assign each location $i$ to the most likely parcel and evaluate the cosine error using the corresponding mean direction:

$$\bar{\epsilon}_{\text{cosine}} = \frac{1}{P} \sum_{i=1}^{P} \left( 1 - \mathbf{v}_{\underset{k}{\text{argmax}}}^{\intercal} \frac{\mathbf{y}_i}{\|\mathbf{y}_i\|} \right), \tag{35}$$

where $\|\mathbf{y}_i\|$ denotes the magnitude of the observed data at location $i$, and $\mathbf{v}_k$ represents the mean direction associated with parcel $k$. The cosine distance is then averaged across all $P$ locations for a given individual.

A more principled formulation leverages the full probabilistic parcellation by computing the expected cosine error under the posterior distribution $\langle \mathbf{u}_i(k) \rangle_q$:

$$\langle \bar{\epsilon}_{cosine} \rangle_q = \frac{1}{P} \sum_{i=1}^{P} \sum_{k} \langle \mathbf{u}_i(k) \rangle_q \left( 1 - \mathbf{v}_k^{\intercal} \frac{\mathbf{y}_i}{\|\mathbf{y}_i\|} \right), \tag{36}$$

where $\langle \mathbf{u}_i(k) \rangle_q$ denotes the inferred posterior probability that location $i$ belongs to parcel $k$.

A limitation of cosine-based measures is that normalization by $\|\mathbf{y}_i\|$ causes locations with low signal-to-noise ratio to contribute equally to the error as those with strong signals. To address this, we introduce a weighting scheme based on the squared magnitude of the data vector, leading to the mean adjusted expected cosine error:

$$\langle \bar{\epsilon}_{\text{Acosine}} \rangle_q = \frac{1}{\sum_i^P \|\mathbf{y}_i\|^2} \sum_i \sum_k \langle \mathbf{u}_i(k) \rangle_q (\|\mathbf{y}_i\|^2 - \mathbf{v}_k{}^T \mathbf{y}_i \|\mathbf{y}_i\|). \tag{37}$$

This formulation effectively weights each location by its signal magnitude, thereby reducing the influence of noisy observations. The resulting metric corresponds to a weighed cosine reconstruction error averaged across all $P$ locations for a given individual, and serves as one of the evaluation metrics in the simulation.

##### S11.2 Proof of the equivalence between the mean adjusted expected cosine error and $1 - R^2$

We provide a mathematical proof that the mean adjusted expected cosine error is proportional to $1-R^2$. Weighting the error by the magnitude of the data vector corresponds to evaluating the squared reconstruction error between $\mathbf{y}_i$ and a prediction scaled to the amplitude of the data, $\mathbf{v}_k\|\mathbf{y}_i\|$. For simplicity, we first consider the hard assignment case and denote $\mathbf{v}_k$ as the predicted mean direction corresponding to the most likely parcel assignment $\mathbf{v}_{\underset{k}{\text{argmax}}}$. The coefficient of determination between $\mathbf{y}_i$ and $\mathbf{v}_k\|\mathbf{y}_i\|$ is defined as:

$$\begin{aligned}
1 - R^2 = \frac{RSS}{TSS} &= \frac{1}{\sum_i \|\mathbf{y}_i\|^2} \sum_i (\mathbf{y}_i - \mathbf{v}_k \|\mathbf{y}_i\|)^2 \\
&= \frac{1}{\sum_i \|\mathbf{y}_i\|^2} \sum_i (\mathbf{y}_i^{\intercal} \mathbf{y}_i - 2\mathbf{y}_i^{\intercal} \mathbf{v}_k \|\mathbf{y}_i\| + \mathbf{v}_k^{\intercal} \mathbf{v}_k \|\mathbf{y}_i\|^2) \\
&= \frac{1}{\sum_i \|\mathbf{y}_i\|^2} \sum_i (\|\mathbf{y}_i\|^2 - 2\mathbf{y}_i^{\intercal} \mathbf{v}_k \|\mathbf{y}_i\| + \|\mathbf{y}_i\|^2) \\
&= \frac{2}{\sum_i \|\mathbf{y}_i\|^2} \sum_i (\|\mathbf{y}_i\|^2 - \mathbf{y}_i^{\intercal} \mathbf{v}_k \|\mathbf{y}_i\|).
\end{aligned} \tag{38}$$

Comparing with Eq. 34, this shows that $1 - R^2 = 2\bar{\epsilon}_{\text{Acosine}}$. Extending this result to the probabilistic setting, where predictions are averaged over parcel assignments using $\langle \mathbf{u}_i(k) \rangle_q$, we have:

$$\langle \bar{\epsilon}_{\text{MSE}} \rangle_q = \frac{1}{\sum_i \|\mathbf{y}_i\|^2} \sum_i \sum_k \langle \mathbf{u}_i(k) \rangle_q (\mathbf{y}_i - \mathbf{v}_k ||\mathbf{y}_i||)^2 = 2\langle \bar{\epsilon}_{\text{Acosine}} \rangle_q, \tag{39}$$

where $\langle \mathbf{u}_i(k) \rangle_q$ denotes the posterior probability of location $i$ belonging to parcel $k$. Thus, the mean adjusted expected cosine error is proportional to the normalized mean squared reconstruction error and is equivalent (up to a factor of 2) to computing $1 - R^2$.

#### Supplemental Figures

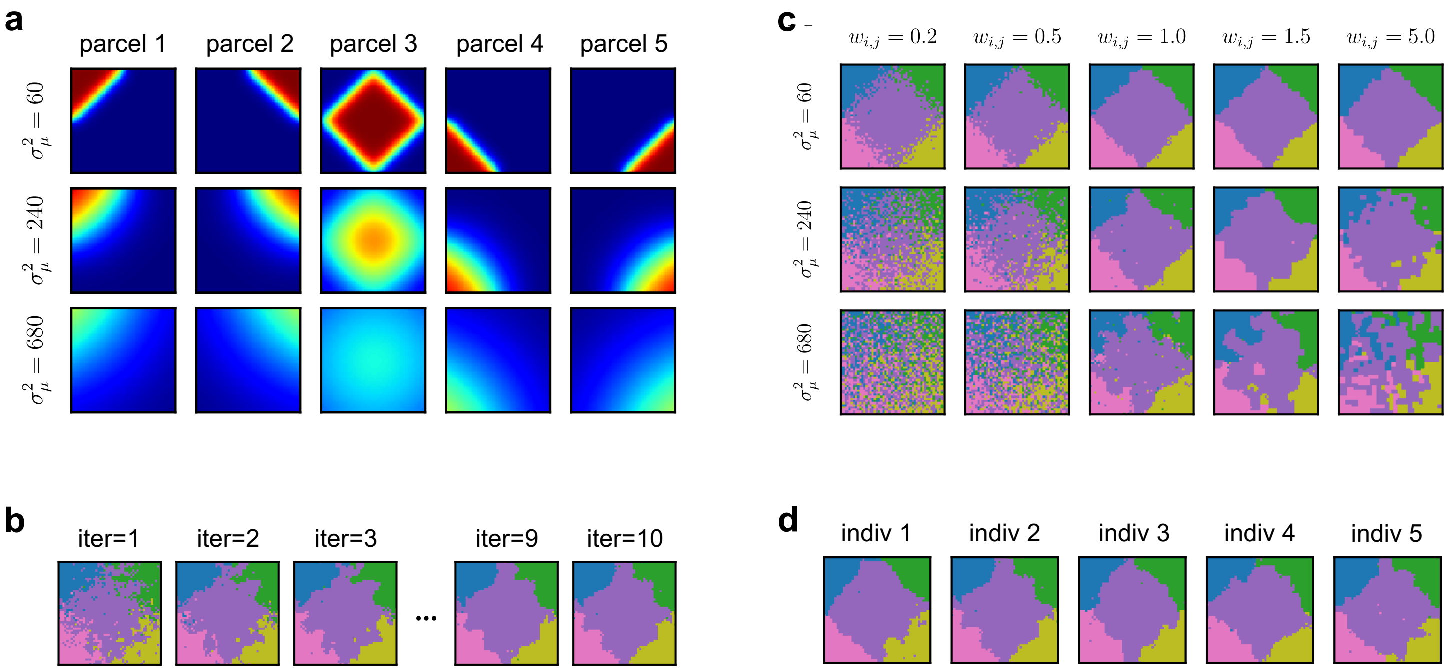


**Supplementary Figure 1: Generation of synthetic datasets. a,** Group-level priors for the five parcels, generated using spatial kernels with varying degrees of smoothness. **b,** Burn-in process used to generate individual-level parcellations. **c,** Example ground-truth individual-level parcellation maps under different parameter settings. **d,** Example ground-truth parcellation maps from different individuals.

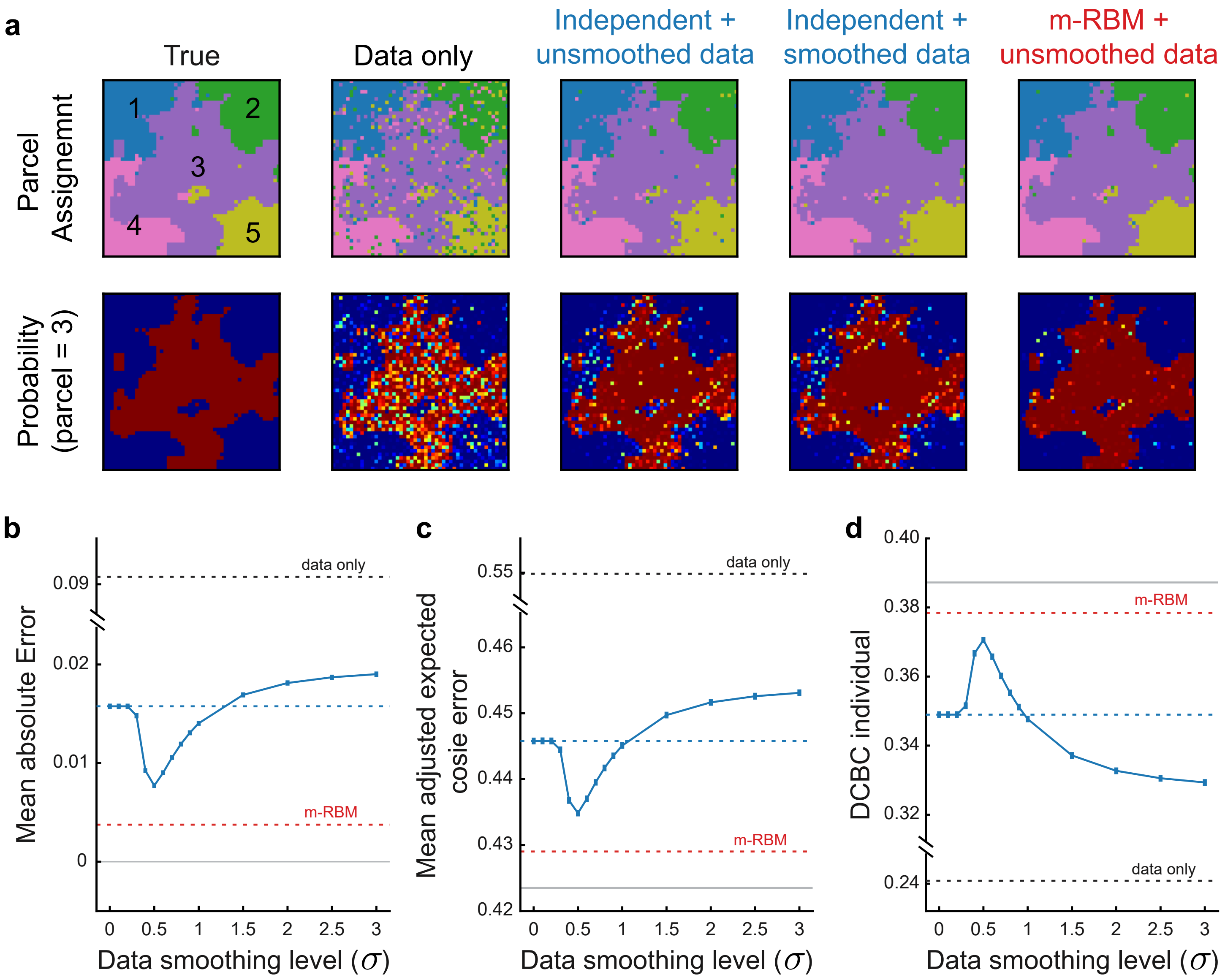


**Supplementary Figure 2: Simulation results. a,** Comparison of reconstructed individual-level parcellations (top) and corresponding probability maps for parcel 3 (bottom), inferred using individual data alone (Data only), spatially independent models with and without smoothing (Independent + unsmoothed data; Independent + smoothed data), and the mRBM model. **b-d,** The mean absolute error (lower is better), mean adjusted expected cosine error (lower is better), and DCDC (Distance-Controlled Boundary Coefficient; higher is better) for individual-level parcellations inferred using the spatially independent model with smoothing (blue solid line) and without smoothing (blue dashed line), and the mRBM model applied to unsmoothed data (red dashed line). The data-only approach (black dashed line) and theoretical optimum (gray solid line) are shown for reference. Error bars indicate the standard error of the mean across 100 simulation replicates.

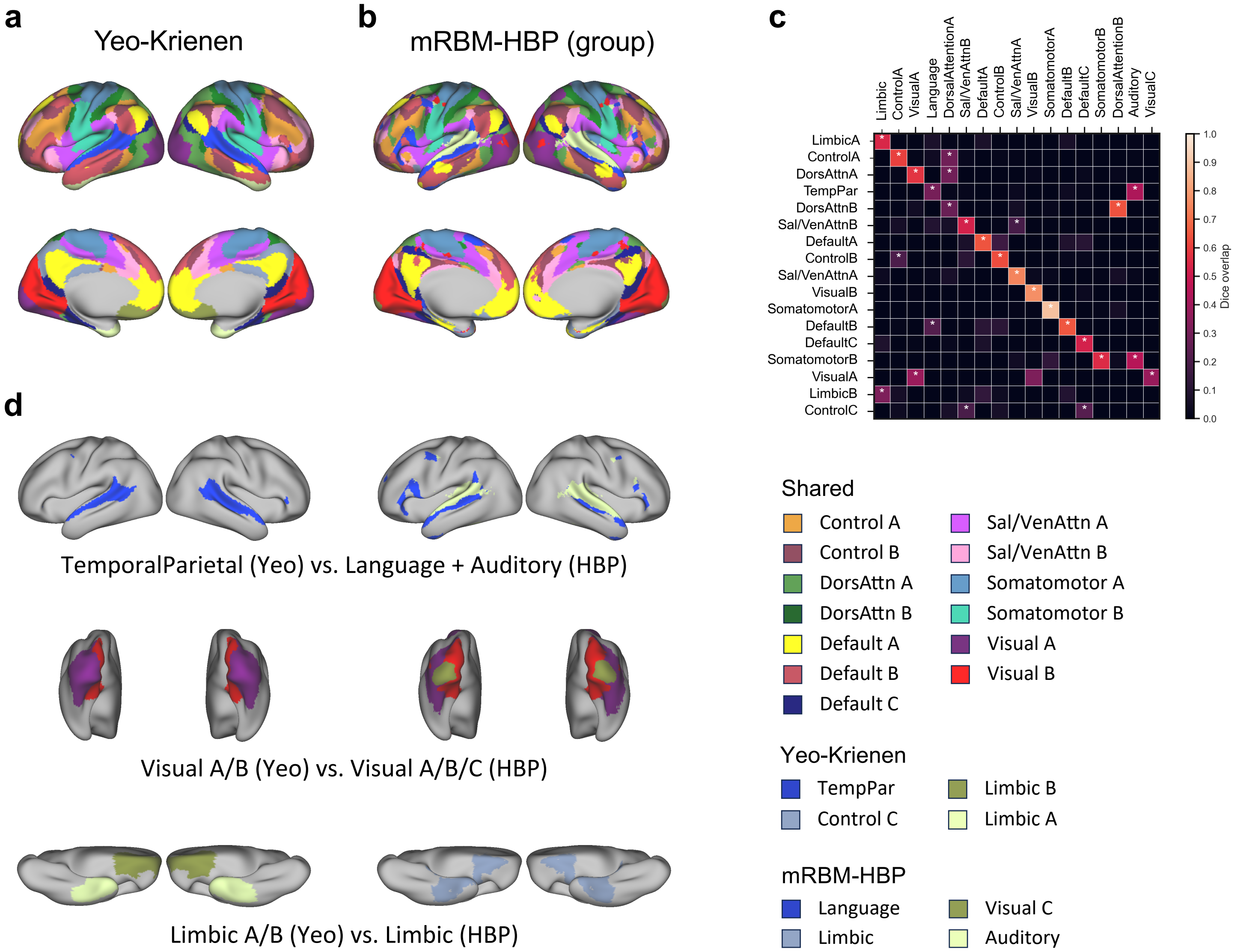


**Supplementary Figure 3: Comparison of the canonical Yeo-Krienen 17-network parcellation and the mRBM-HBP group-level resting-state parcellation. a,** Canonical Yeo-Krienen 17-network cortical parcellation. **b,** mRBM-HBP 17-network resting-state group-level parcellation. **c,** Dice overlap between the Yeo-Krienen and mRBM-HBP parcellations. **d,** Functional networks showing major differences between the two parcellations.

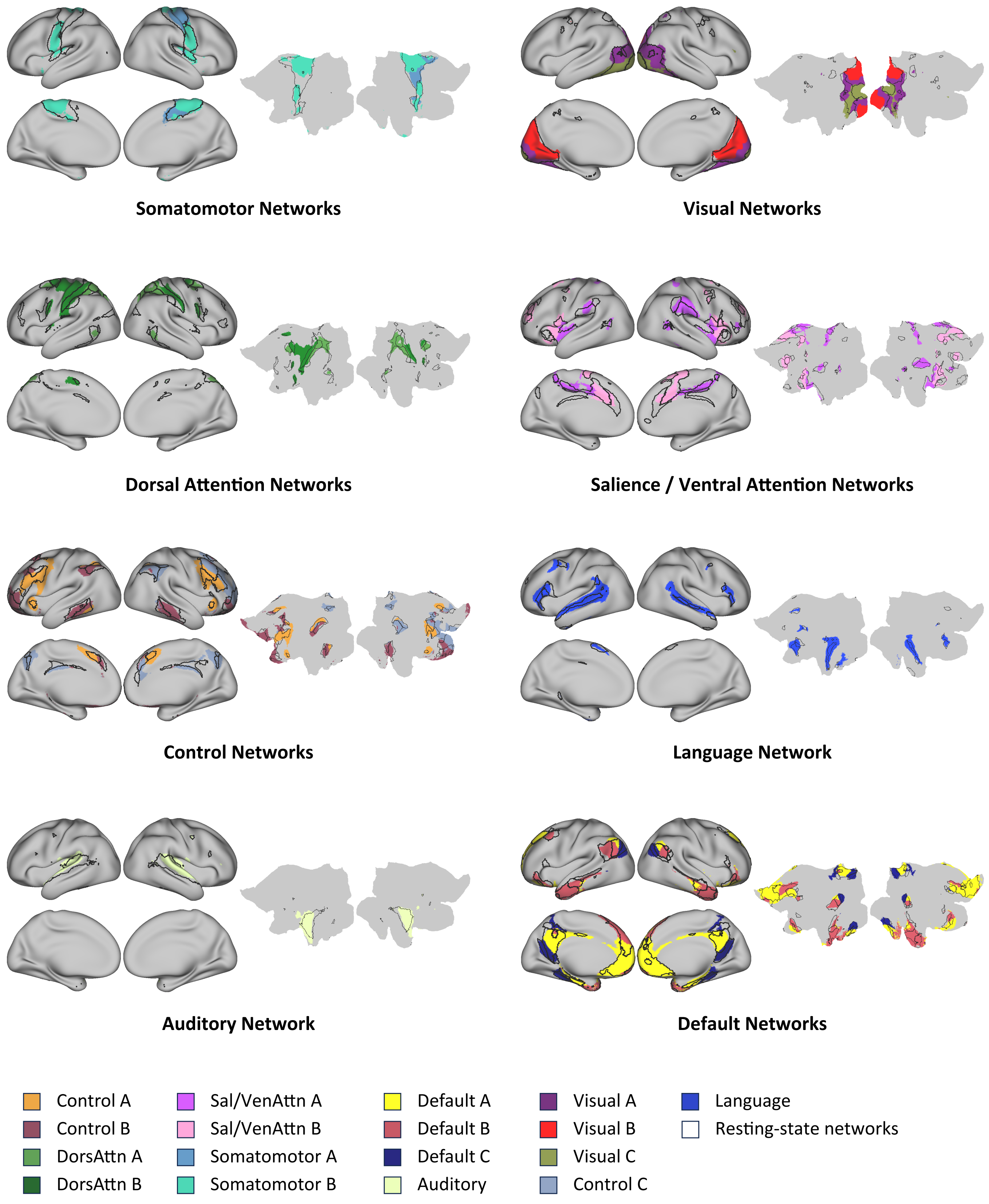


**Supplementary Figure 4: Network-level comparison of resting-state and task-based 17-network group-level cortical parcellations generated by mRBM-HBP.** Task-based parcellations are shown in color, with corresponding network boundaries derived from the resting-state parcellation overlaid.

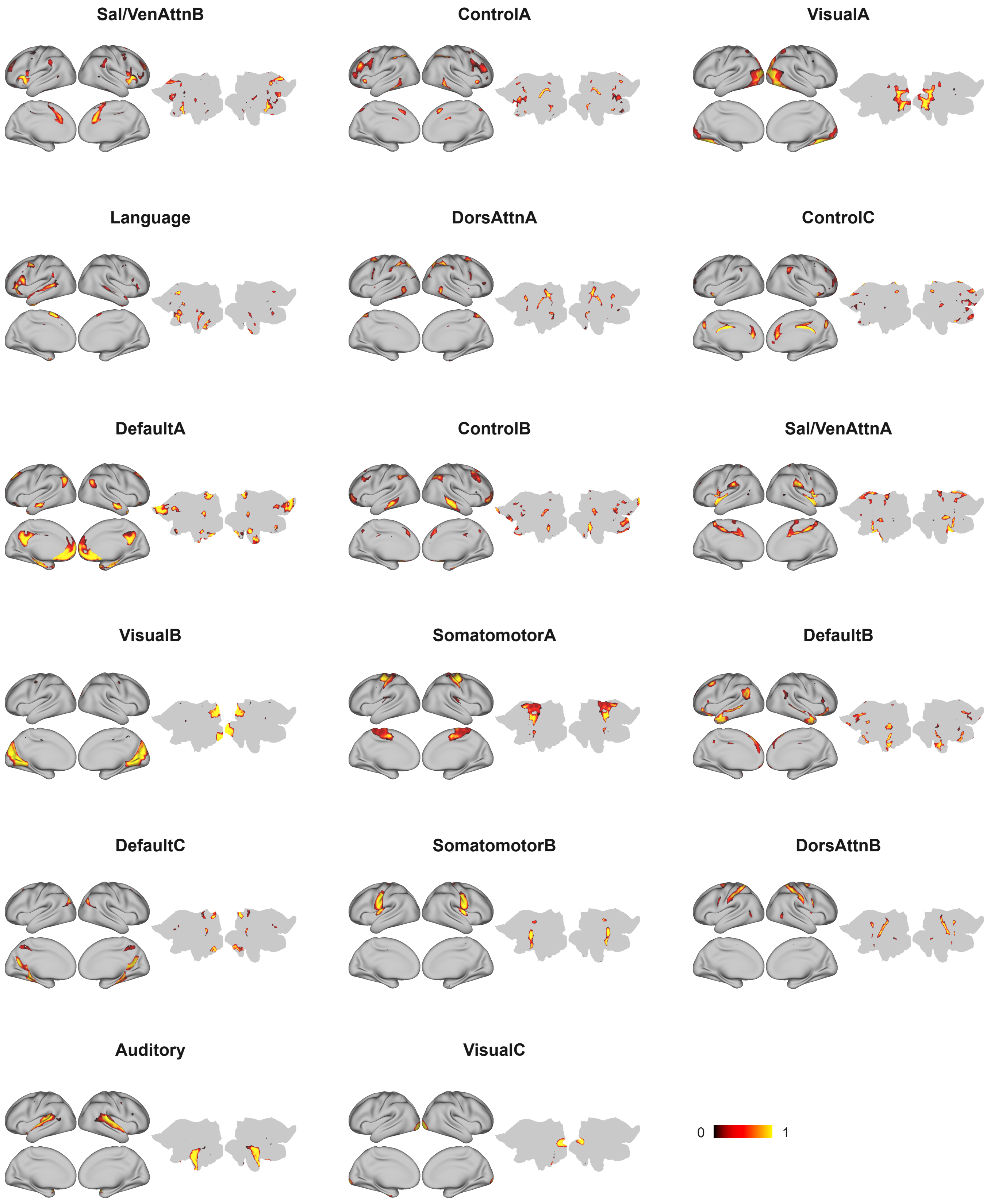


**Supplementary Figure 5: Network-wise probability maps for the fused task-rest 17-network group-level parcellation.** Color maps show the full probability distribution for each network using a consistent scale across networks. For visualization, probability maps are masked by their corresponding binary network assignments; therefore, displayed values may not start at zero.

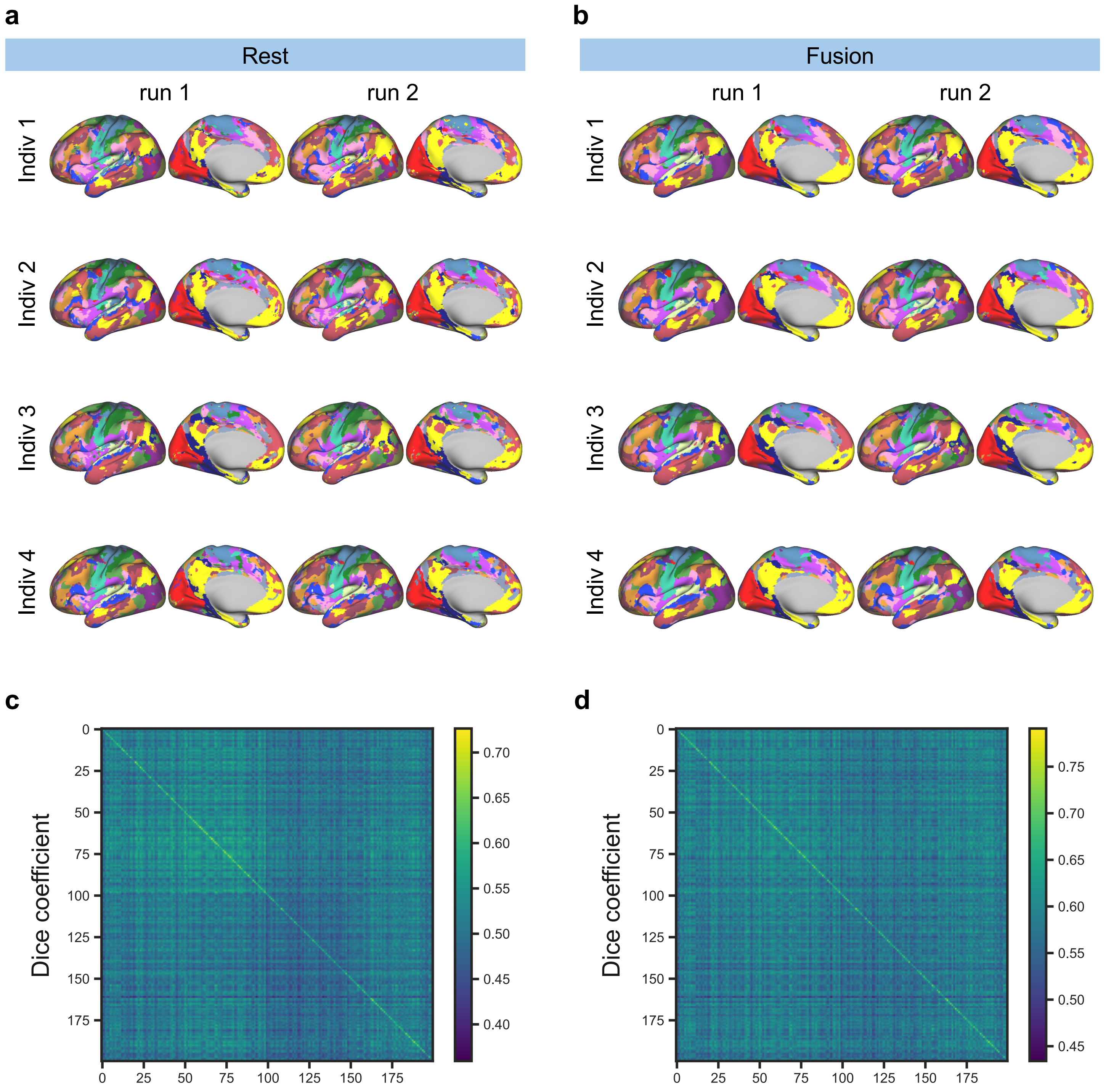


**Supplementary Figure 6: Within-individual similarity and between-individual variability of parcellations in HCP-YA. a-b,** Example individual-level parcellations generated using either the resting-state or fused prior from independent runs of the same individual. **c-d,** Dice similarity of parcellations generated using either the resting-state or fused prior, computed between independent runs from the same individual to quantify within-individual similarity, or between pairs of different individuals to quantify between-individual similarity.

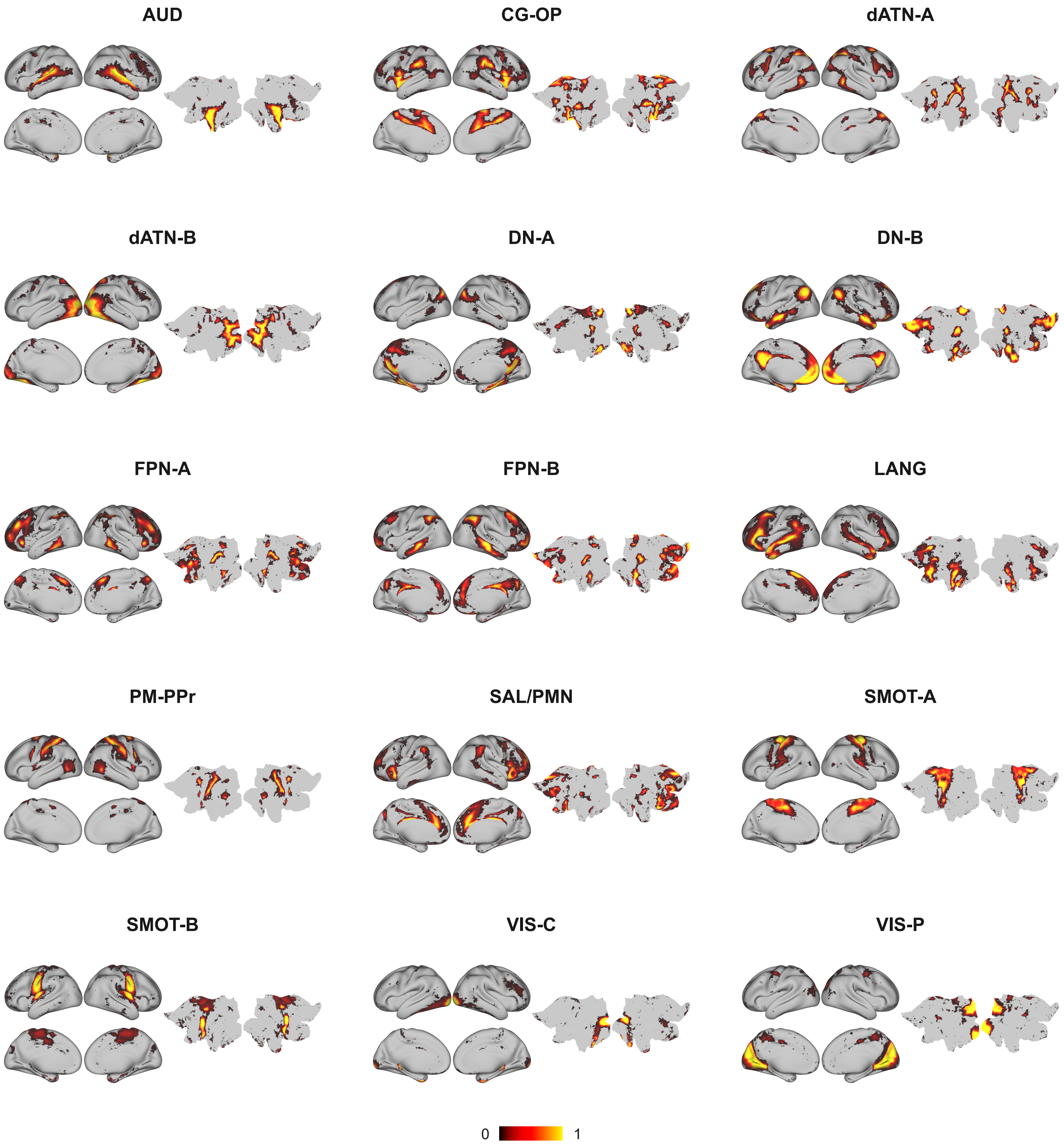


**Supplementary Figure 7: Network-wise probability maps for the fused task-rest 15-network group-level parcellation.** Color maps depict the full probability distribution for each network using a consistent scale across networks. For visualization, probability maps are masked by their corresponding binary network assignments; therefore, displayed values may not start at zero.

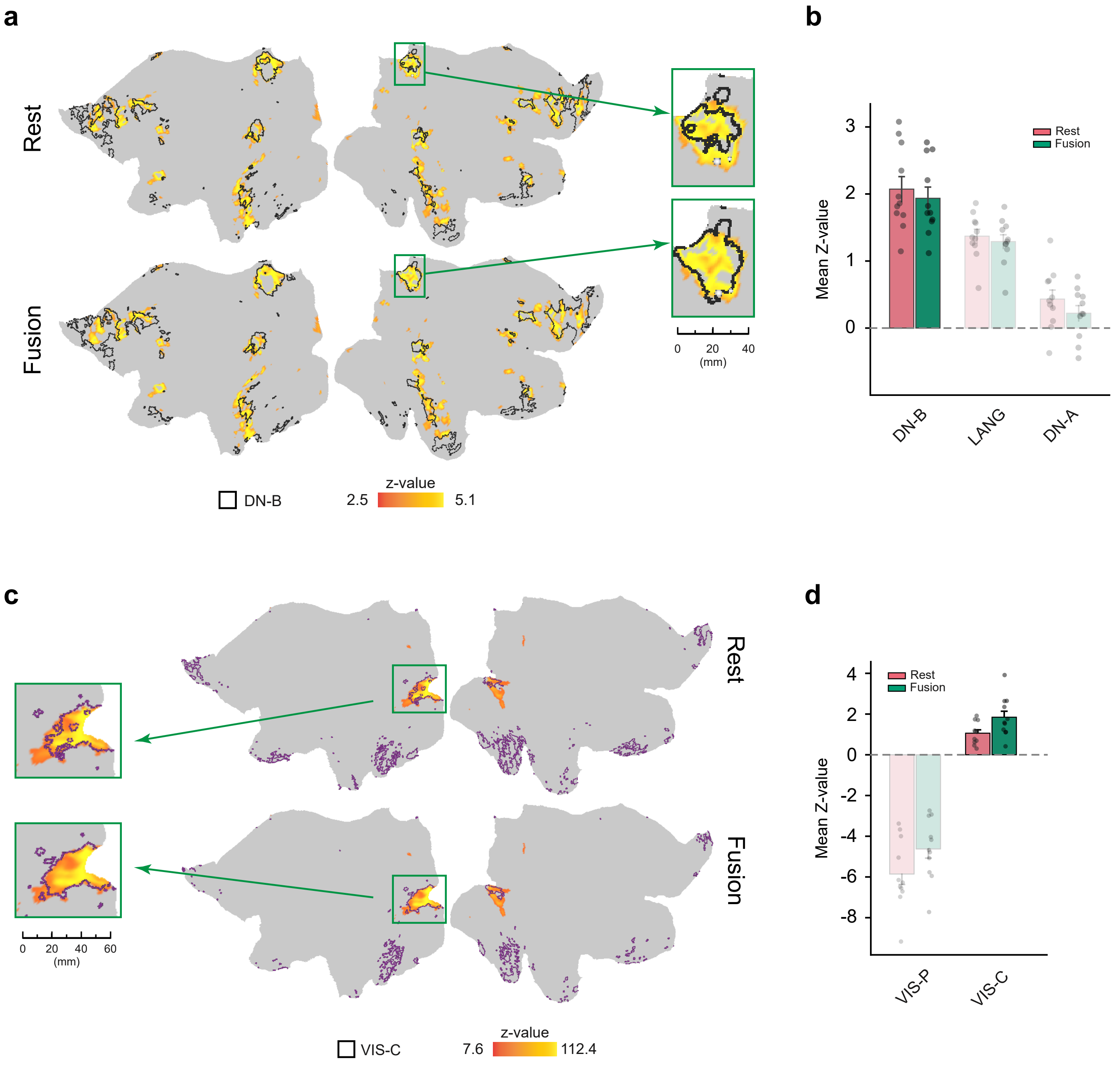


**Supplementary Figure 8: Additional examples of precision functional mapping in HPN. a,** Thresholded contrast map from the theory-of-mind (ToM) task for an HPN participant, displayed on the flattened cortical surface and overlaid with DN-B network boundaries derived from the resting-state and fused priors using a single resting-state run. **b,** Average ToM contrast z-values within DN-B and adjacent networks, DN-A and LANG, defined using the resting-state and fused priors across HPN participants. **c,** Thresholded contrast map from the visual task for an HPN participant, displayed on the flattened cortical surface and overlaid with VIS-C network boundaries derived from the resting-state and fused priors using a single resting-state run. **d,** Average visual contrast z-values within VIS-C and the adjacent VIS-P network, defined using the resting-state and fused priors across HPN participants. In **b** and **d**, error bars represent the standard error of the mean.

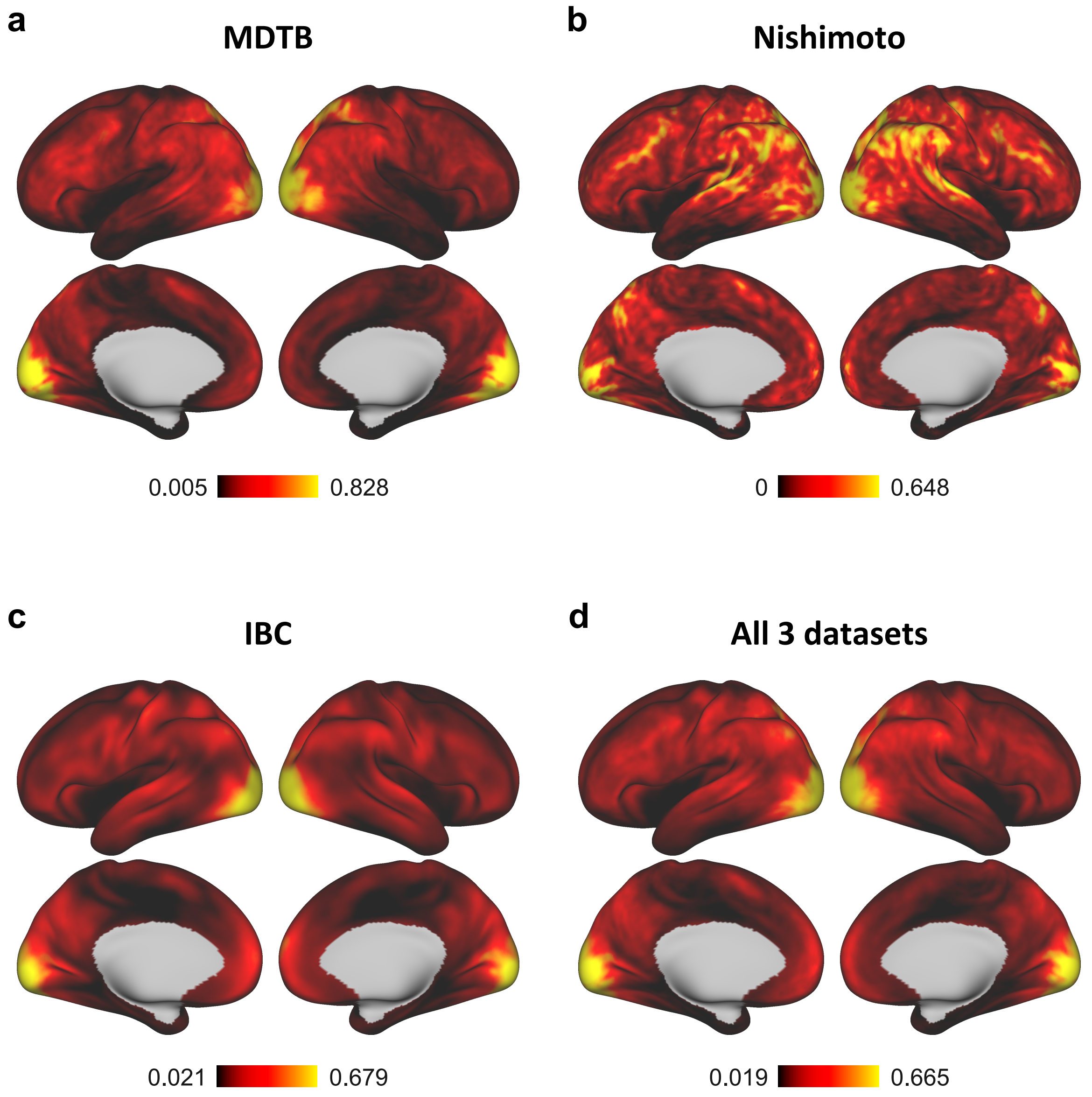


**Supplementary Figure 9: Converge maps of task activation profiles across datasets. a-c,** Coverage maps showing the frequency with which each vertex was included among the top 10% most activated or bottom 10% most deactivated vertices across task activation maps in the MDTB, Nakai & Nishimoto, and IBC datasets. **d,** Combined coverage map across all three datasets.
